# Highly efficient genotype compression leveraging genealogical relatedness

**DOI:** 10.64898/2026.04.29.721594

**Authors:** Amber Shen, Xinran Wang, Nicholas Mancuso, Luke J. O’Connor

## Abstract

Large genetic datasets are terabytes in size, presenting a computational challenge that will intensify as sequencing efforts scale. We present a lossless compression algorithm, *kodama*, which supports matrix multiplication and is suitable for large-scale statistical analyses. *Kodama* leverages genealogical relatedness among nominally unrelated individuals and infers a novel data structure similar to the ancestral recombination graph (ARG), called the linear ARG. We applied *kodama* to whole genome sequencing data from UK Biobank and All of Us. Inferred linear ARGs were 17-89 times smaller on disk compared to the input data; the entire UK Biobank N=200k dataset can be loaded into memory (58GB). Compared with the recently proposed genotype representation graph (GRG), the linear ARG is 2.5 times smaller. Genotype matrix multiplications, which are the bottleneck in most statistical applications, are extremely fast with the linear ARG; we performed a GWAS on the UK Biobank 200k cohort across 89 traits with 42 covariates in 100 seconds, representing a 4,700-fold speedup over PLINK 2.0. We expect that the linear ARG will enable genetic analyses to scale to millions of samples.

## Introduction

Millions of human genomes have been sequenced or genotyped, presenting an opportunity for genetic discovery but also substantial computational and storage challenges^1^. Whole genome sequencing (WGS) datasets can be terabytes in size, and modern biobanks pair large-scale WGS data with rich phenotypic data, like electronic healthcare records and medical images, making comprehensive analyses expensive^2,3^. REGENIE, a state-of-the-art genetic association method optimized for speed, takes around 60 CPU hours to run the linear regression step for a single trait GWAS on the UK Biobank 500k cohort^4^. Biobanks continue to grow; Our Future Health plans to genotype 5 million individuals^5^, and the computational cost of analyzing such a dataset could cause potential discoveries to be missed.

A possible solution is to leverage the genetic relatedness of study participants. As sample size grows, each added sample has closer genealogical relatives who are already in the study, such that nonredundant genetic information grows sublinearly^6^. A compression algorithm that leverages genealogical structure could make large datasets dramatically smaller, with no tradeoff in accuracy^7^. Such a method would be especially useful if the mathematical operations used in statistical applications could be applied to the compressed data structure directly, without decompressing it first^8,9^. Of particular importance is the genotype matrix-vector product *X*′*y*, which is a rate-limiting step in genome-wide association studies (GWAS).

One existing compression algorithm is the Positional Burrows-Wheeler Transform (PBWT), which leverages haplotype structure by compressing identical subsequences^10^. The PBWT data structure supports efficient algorithms for sequence matching, making it ideally suited for reference-based genotype imputation and phasing^11,12^, and it is the basis of several compressed file formats, like Savvy, BGT, and XSI^13–15^. However, the PBWT is not ideally suited for statistical applications, like GWAS, that rely on genotype matrix operations.

A different approach is to infer the genealogical process itself, in the form of an ancestral recombination graph (ARG)^16,17^. ARGs can be inferred from genetic data^7,18–22^, and ARG inference is an attractive approach for genealogical compression because the parsimony of the genealogical process is the reason that compression is possible. Simulations suggest that an ARG for the entire human population could be just one terabyte in size, vs. petabytes for a traditional VCF^7^. Compressing real datasets remains a challenge; for example, *tsinfer* often makes real datasets larger rather than smaller^7^. Recently, the *threads* method achieved a compression ratio of 14x on a UK Biobank imputed dataset^23^ and has been used for accelerated matrix multiplication^24^. Similarly, a recently proposed ARG-like data structure, the genotype representation graph (GRG), compresses biobank-scale WGS data by ∼13x; this method is expressly designed for matrix multiplication and is several times faster than a conventional approach^9^.

We make three contributions. First, we introduce an ARG-like data structure, the *linear ARG*, whose vertices are linear combinations of their ancestors. The linear ARG is related to the genotype matrix by an algebraic formula, making it well suited for statistical applications. Second, we introduce an inference method, *kodama*, which achieves state-of-the-art compression on biobank-scale datasets. Third, we implement highly efficient programs for matrix-vector multiplication, and we develop a fully featured method for genetic association testing with significant computational advantages over conventional methods.

### Overview of methods

Here, we present a high level description of our data structure, with details provided in Supplementary Note. At a locus without recombination or recurrent mutation, there exists a genealogical tree such that every variant can be mapped to a single edge (*j, i*) of that tree; node *i* is the “mutation node”, and the carriers of the derived allele are the descendants of *i*. This representation requires less memory than the corresponding genotype matrix *X*, while encoding exactly the same data (Figure 1a). Moreover, if *y* is some vector of phenotypes for individuals encoded in this tree, then the genotype matrix product *X′y* can be computed by traversing the tree. To see this, the traversal is initialized by assigning to leaf nodes, which represent samples, corresponding values from the input vector *y*. The tree is traversed from bottom to top, and each internal node is assigned the sum of its child nodes (Figure 1b). Entry *i* of *X′y* is the value assigned to the mutation node of variant *i*. The number of addition operations is the number of edges in the tree, which can be much smaller than the size of *X*.

**Figure 1:**
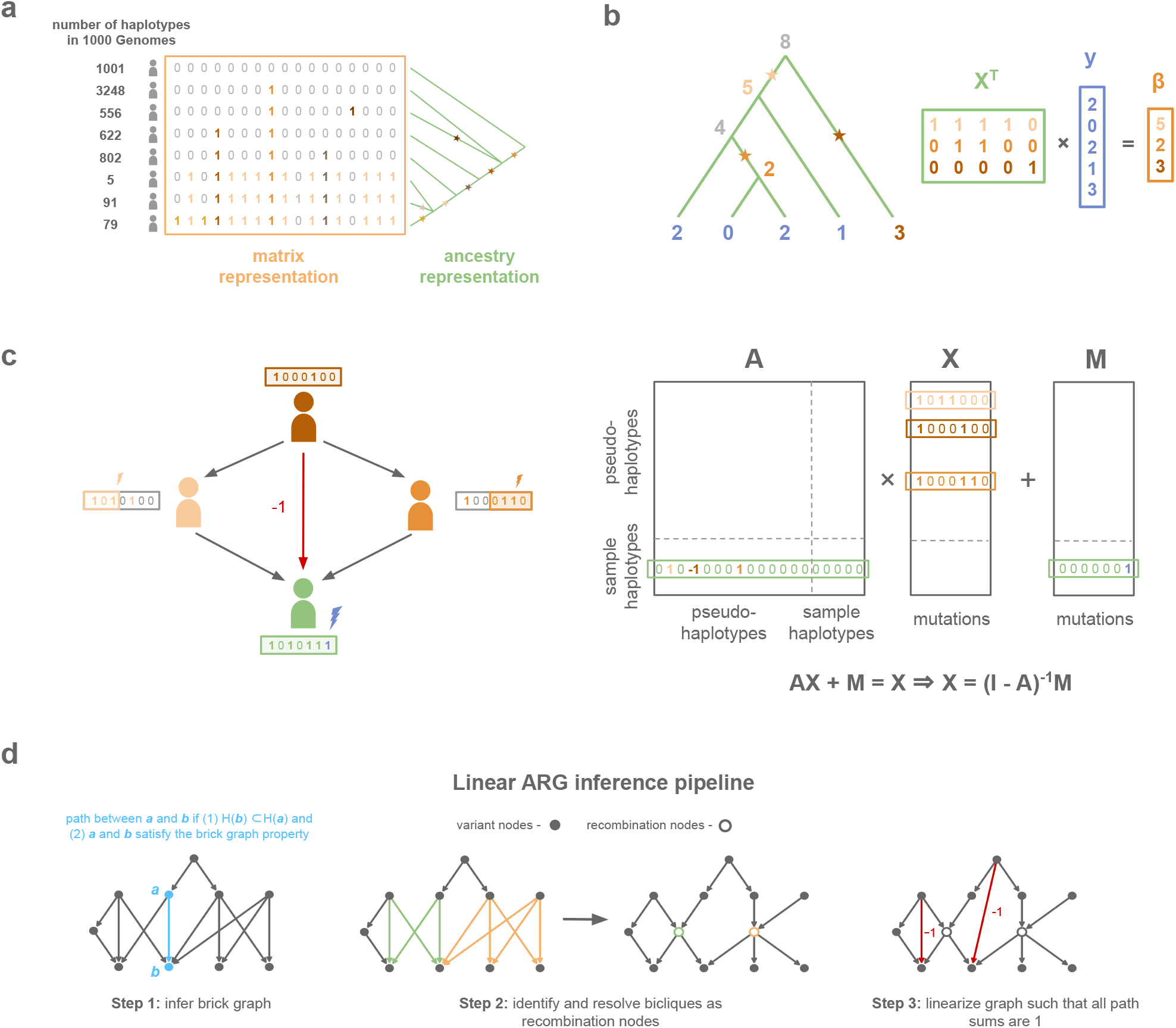
Overview of the linear ARG compression scheme. **a**, Comparison of the matrix representation and ancestry representation for chr2:234,383,207-234,384,556 in 1000 Genomes^25^. The matrix representation stores each variant as many times as there are carriers while the ancestry representation only needs to store the position of the variant on the tree. **b**, Toy example of matrix multiplication with the genotype matrix using a graph traversal on the genealogical tree in the case of no recombination. **c**, The sample haplotype (green) is a composite of its ancestral pseudo-haplotypes (orange) and *de novo* mutation (purple). **d**, The linear ARG can be inferred from a phased genotype matrix in three steps: (1) brick graph inference, (2) find recombinations, (3) linearization.

This procedure corresponds to an algebraic equation. Let node *i* be the child of node *j*, with variants indicated by *m*_*i*_, then its haplotype is given by *x*_*i*_ = *x*_*j*_ + *m*_*i*_. Concatenating the *N* vectors *x* into a matrix *X*, and the *N* vectors *m* into a matrix *M*, this equation becomes

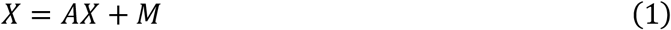

where *A* is the adjacency matrix, with *A*_*ij*_ = 1 if haplotype *X*_*i*_ is the child of haplotype *X*_*j*_. This equation can be solved for *X* as,

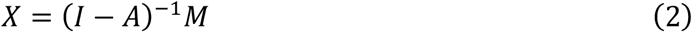

The right-hand side of (2) can be substituted into any algebraic expression involving *X*. In particular, the expression *X′y* becomes *M*′(*I* – *A*′)^−1^*y*, and evaluating this expression involves solving a sparse triangular system of equations, since *A* is a directed acyclic graph. This procedure is exactly the graph traversal described above. The rows of *X* include both sample haplotypes, belonging to individuals in the study, and ancestral haplotypes, representing inherited sequences; the genotype matrix of sample nodes only is given by the expression *S*(*I* − *A*)^−1^*M*, where *S* contains one non-zero entry per sample.

In the presence of recombination, a haplotype can have multiple parents, and each parent transmits alleles at some sites but not others (Figure 1c). As a result, haplotypes are not linear combinations of their ancestors in the ARG. We define a modified ARG, called the *linear ARG*, which restores linearity. The linear ARG differs from an ordinary ARG in two ways. First, in the linear ARG, a node has the same descendants at every position, such that alleles are always transmitted. Nowbandegani & Wohns et al. called such nodes “bricks”^26^. Second, the edges of the linear ARG have integer-valued weights, and in particular, certain edges have a weight of -1. For example, if a node *i* has two parents *j, k*, and those parents themselves have a common ancestor *l*, then we see,

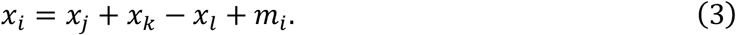

Any alleles carried both by *j* and by *k* are double counted in *x*_*j*_ + *x*_*k*_, but then cancelled out by the negative term −*x*_*l*_. The expression *x*_*j*_ + *x*_*k*_ – *x*_*l*_ corresponds to row *i* of *A*, which has entries 1,1,-1 in positions *j, k, l*. Negative edges are needed to allow “diamond-shaped” paths in the graph, the existence of which distinguishes the linear ARG from the GRG^9^. Any ARG corresponds to a single linear ARG, and we show this correspondence formally in the Supplementary Note.

We developed a method, *kodama*, to infer the linear ARG. It proceeds in three steps. First, it generates an initial graph, the *empirical brick graph*, with one node per mutation (Figure 1d). In this graph, a path connects derived allele *a* with derived allele *b* if the carriers of *b* are a subset of the carriers of *a*, and if an additional condition is satisfied (Supplementary Note). We compute the empirical brick graph using an algorithm, the brick graph algorithm, which iterates twice through the genome, once forward and once backward (Supplementary Note). In the absence of recombination, the empirical brick graph is a genealogical tree, and more generally, the empirical brick graph alone is useful for compression when the recombination rate is low. However, when the recombination rate is high, the empirical brick graph contains inefficient ‘bicliques’ due to recombination events which are not tagged by any single mutation (Figure 1d). In the second step of *kodama*, we detect these bicliques and factor them to produce a brick graph with more nodes but fewer edges (Figure 1d). In the third step, to linearize this graph, we add weighted edges to the graph such that for all nodes that have a path between them, the sum of their path weights is one. This step is not needed for compression, but it is needed for the weighted adjacency matrix to satisfy equation (2) (Figure 1d). We provide details of each step in the Supplementary Note and specify the format of the linear ARG data structure in the Methods section.

### Genealogical compression

To assess the data compression performance of our approach, we applied *kodama* to phased whole genome sequencing data (N=200k) from UK Biobank^27,28^ for chromosomes 1 through 22, partitioning the data into roughly 20Mb blocks (Methods). We verified that the compression was lossless by confirming the linear ARG allele counts matched the allele counts from the genotype matrix (Methods). We applied GRG and XSI to the same data, for chromosomes 1, 11, and 21; these methods were chosen because they support matrix multiplication (see below). We quantified the size on disk of the resulting data structures as compared with the original vcf.gz files, as well as the PLINK 2.0 pgen file format, which leverages haplotype structure for compression by storing differences between columns of the genotype matrix^29^. We also quantified size in memory for our linear ARGs, the GRG, and a generic sparse genotype matrix format (compressed sparse column format, CSC; the adjacency matrix of the linear ARG is formatted similarly).

The linear ARG (27GB) was 89 times smaller than the input .vcf.gz file on disk. In memory, the linear ARG was 58GB, small enough that the entire UK Biobank N=200k dataset can be loaded into computer memory (Table 1); it was 58 times smaller than a sparse genotype matrix with one entry per minor allele, and 2.5 times smaller than the GRG data structure (Figure 2a). On disk, compared to other haplotype-informed compression methods, the linear ARG was 2.5 times smaller than GRG, 15 times smaller than pgen, and 3 times smaller than XSI (Figure 2b). The improvement over XSI is expected to be representative of how linear ARGs would compare with other methods based on the PBWT, like Savvy and BGT; DeHaas et al. found that XSI and Savvy have similar compression ratios^9^. We did not compare with *tsinfer*, which achieves very large compression ratios in simulated data but not in real data^7^.

**Table 1:** Compression performance. Disk sizes before and after compression and size in memory after compression. NNZ (number of non-zeros) ratio is the number of minor alleles in the genotype matrix over the number of non-zero entries in the linear ARG. Linear ARG size in memory and on disk excludes metadata; vcf.gz size on disk includes it. *We filtered multi-allelic sites while streaming in AoU data to minimize disk transfers. This GB size reflects original unfiltered size. The reported number of variants is after filtering.

| Dataset | Number of samples | Number of variants | NNZ ratio | Linear ARG size in memory (GB) | Linear ARG size on disk (GB) | Linear ARG metadata size on disk (GB) | vcf.gz size on disk (GB) |
| --- | --- | --- | --- | --- | --- | --- | --- |
| 1000 Genomes | 6,404 | 70,629,015 | 15.9 | 5.7 | 2.1 | 0.5 | 35.0 |
| UK Biobank 200k | 199,911 | 659,887,507 | 75.6 | 58.0 | 26.5 | 3.5 | 2347.1 |
| UK Biobank 200k<br>MAF > 0.01 | 199,911 | 9,886,035 | 130.7 | 30.1 | 12.9 | 0.09 | 527.6 |
| UK Biobank 200k<br>chr 1, 11, 21 | 199,911 | 93,742,553 | 75.5 | 8.2 | 3.7 | 0.5 | 333.7 |
| UK Biobank 200k chr 21 | 199,911 | 8,463,891 | 70.0 | 0.86 | 0.39 | 0.04 | 30.7 |
| UK Biobank 500k chr 21 | 490,541 | 9,479,292 | 72.7 | 2.16 | 1.0 | 0.1 | 49.5 |
| All of Us chr 21 (v8) | 414,830 | 6,185,871 | 74.1 | 1.05 | 0.49 | 0.06 | 40.9* |

**Figure 2:**
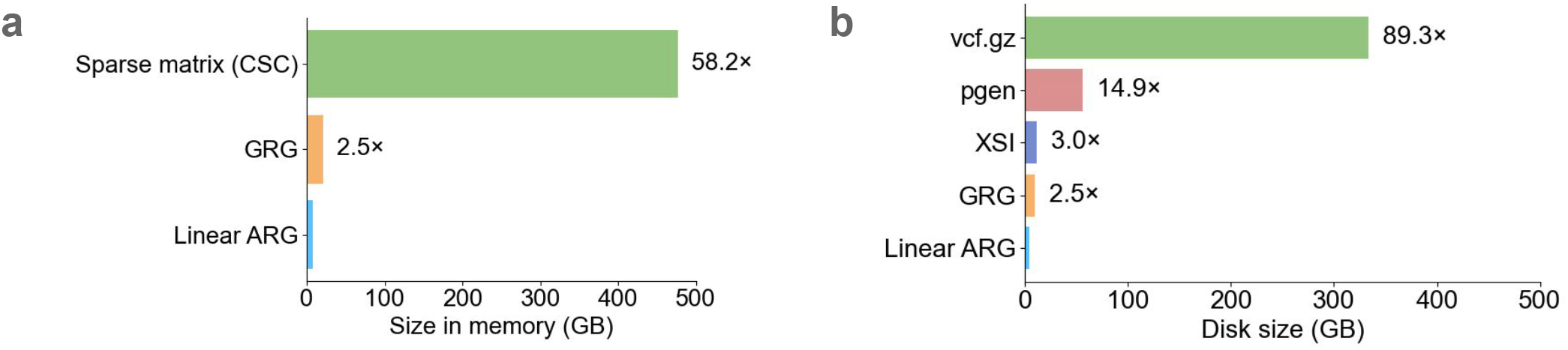
Comparison of compression methods. Methods were evaluated on chromosomes 1, 11, and 21 of UK Biobank (N=200k). **a**, Size in memory for the linear ARG, genotype representation graph (GRG), and compressed sparse column matrix (CSC). **b**, size on disk for the input data (vcf.gz), PLINK 2.0 compressed format (pgen), a format based on the Positional Burrows-Wheeler transform (XSI), GRG, and linear ARG. For numerical results, see Supplementary Tables 1, 2, and 5-7.

Many statistical applications involve common variants only. We inferred common-variant linear ARGs (MAF>0.01) (Methods); these are 30 GB in memory (vs. 58 GB with all variants) (Table 1), and they accelerate matrix multiplication significantly (see below). In general, rare variants are less compressible than common variants, but also much smaller before being compressed. For example, a singleton variant can be encoded by simply recording the haplotype which carries it, and further compression is impossible but also not necessary. In UK Biobank (N=200k, chromosomes 1, 11, 21), the common variant linear ARG is 110 times smaller than the sparse genotype in memory (Supplementary Table 5).

We inferred linear ARGs on three other datasets: 1000 Genomes^25^ (N=3,202, chromosomes 1-22), All of Us v8^3^ (N=414,830, chromosome 21), and the recently released N=500k phased UK Biobank dataset^2,27^ (chromosome 21) (Table 1) (Methods). On the large All of Us and N=500k UK Biobank (UKB) datasets, the compression ratios were 73 and 74 respectively (vs. 70 for the UK Biobank 200k dataset on chromosome 21). The much smaller 1000 Genomes dataset had an NNZ ratio of 16, matching the intuition that when datasets grow, each added individual has closer genealogical relatives who are in the dataset already, helping compression.

### Simulated genealogies

In a simulated genealogy, the generative ARG is expected to approximate the smallest possible representation of the simulated data. We simulated 100 ARGs using *msprime*^30,31^ under an Out of Africa model^32^ over a 1Mb region of chromosome 21 using stdpopsim^33,34^ (Methods) and inferred corresponding linear ARGs. We compared the number of edges in the inferred linear ARGs to the number of edges in the data-generating ARGs, noting that an optimal linear ARG is expected to have more edges than an optimal ARG (Supplementary Note).

At a sample size of 100,000, the linear ARG had 1.25 times more edges than the generative ARG and 271 times fewer non-zero entries than the genotype matrix (Figure 3a). It performed less well at smaller sample size: with a sample size of 10,000 it had 2 times more edges than the generative ARG and 69 times fewer non-zero entries than the genotype matrix (Figure 3a). Previous studies found that *tsinfer* achieves near-optimal compression in simulated datasets, both at large and at small sample size^7^, and that GRG achieves near-optimal compression only at small sample size^9^ (the opposite of what we observe). We are unsure the reason for these apparent differences. Demography is expected to influence compression differently at different sample sizes; in particular, population expansion should cause compression ratios to plateau in large samples due to the relative depletion of recent relatives.

**Figure 3:**
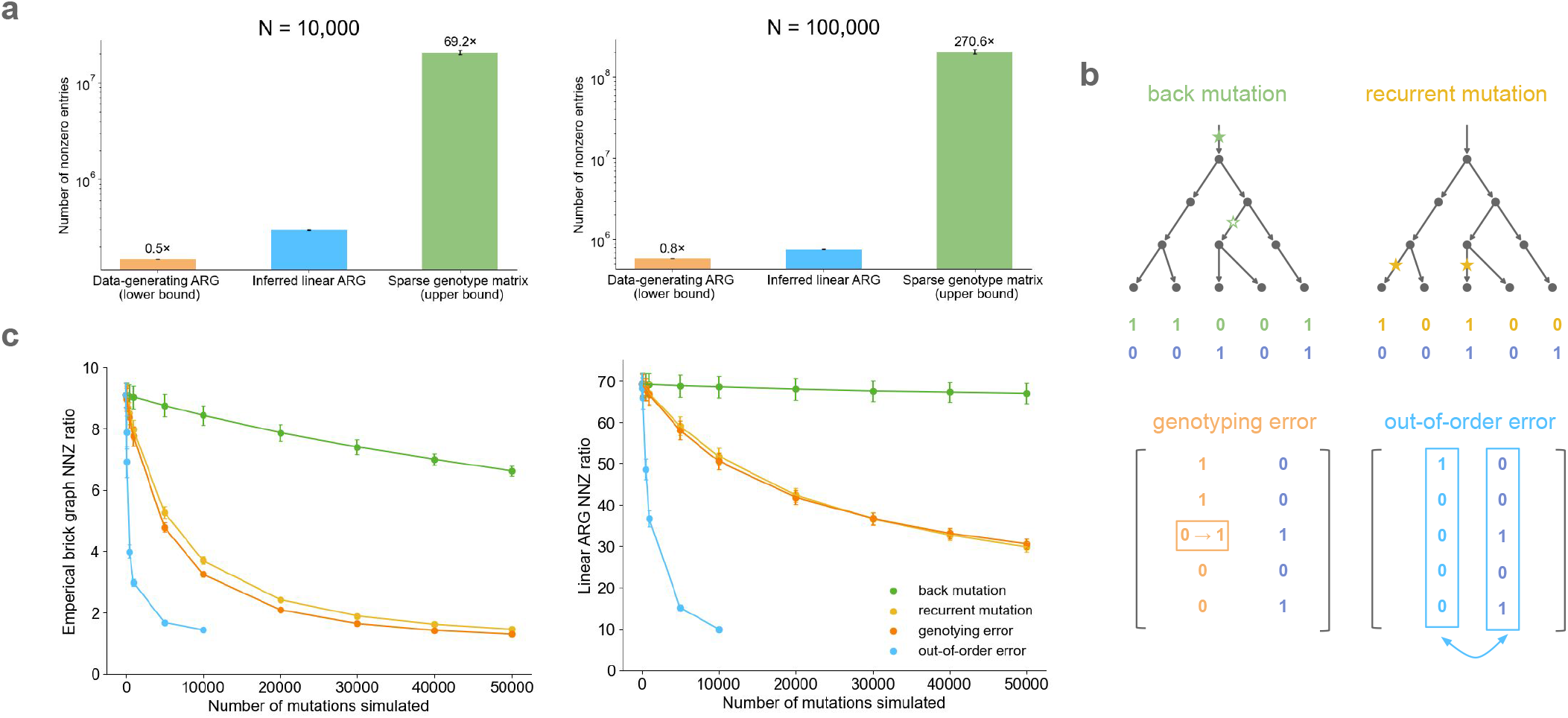
Genealogical compression of simulated genotype data. **a**, Number of non-zero entries (NNZ) between the simulated data-generating ARG (most compressed), the inferred linear ARG, and the sparse genotype matrix (no compression). **b**, Overview of the four kinds of mutations and errors simulated. **c**, The brick graph and linear ARG NNZ ratio (number of minor alleles over number of non-zeros in the linear ARG) versus the number of mutations and errors simulated on top of the *msprime* simulated genotype matrices. For numerical results, see Supplementary Tables 8 and 9.

These simulations assume an infinite-sites model and neglect the influence of genotyping error. Genotyping errors and recurrent mutations affect compression because they produce variants which are inconsistent with their local genealogical tree. Beginning with the same simulated data (N=10,000 haplotypes, M∼63,000 variants), we simulated four kinds of mutations or errors (Figure 3b): recurrent mutations, back mutations, genotyping (allele-flipping) errors, and out-of-order errors (Methods). We simulated up to 50,000 events of each type.

Overall, the compression ratio decreases with the number of mutations or errors, but it does so slowly for three out of the four event types. For back mutation, there was nearly no effect; this is likely because most simulated mutations involved rare variants, such that the two mutations occurred close together in the genealogical tree. For recurrent mutations and genotyping errors, there was a noticeable but modest effect. To understand why the linear ARG was robust to these events, we computed a compression ratio between the number of non-zeros in the genotype matrix and the number of non-zeros in the empirical brick graph (the output of step one in Figure 1d). The empirical brick graph is strongly affected by errors (except for back mutations) (Figure 3c). This indicates that recurrent mutations and genotyping errors resemble recombination events, such that they are rescued by the find-recombination step (step two in Figure 1d).

The fourth event type involved switching the genomic location of two faraway variants. This error type had a dramatic effect on the compression ratio (Figure 3c), probably because it causes the tree to be completely reshuffled instead of causing a local change in the tree topology. In principle, this kind of error could be produced if reads were misaligned to the reference genome; however, this is not expected to occur in most regions of the genome.

### Genome-wide association

We implemented a method, *kodama-gwas*, for linear regression GWAS with the linear ARG (Methods). *kodama-gwas* computes the effect size and standard error for each variant in the linear ARG on one or more phenotypes, with any number of covariates. It parallelizes computations over blocks of the genome, and it is optimized for multi-trait analyses. In its default, fastest configuration, *kodama-gwas* uses two approximations, both relating to the variance of genotype residuals. First, when there are multiple phenotypes, it computes residual variances once and reuses these values across phenotypes (the same approach is used by REGENIE^4^). This calculation is approximate when phenotypes have missing values and missingness differs among phenotypes. It has a dramatic effect on runtime when there are multiple traits and multiple covariates. The ‘—qt-residualize’ option in PLINK 2.0^29^ has a similar effect on runtime but makes a less accurate approximation (see Methods). Second, the default configuration assumes Hardy-Weinberg equilibrium (HWE) conditional upon covariates. This assumption has a smaller effect on runtime, but relaxing it requires a modification to the data structure. This modification allows us to compute the matrix product *Z*′*y* where *Z*_*ij*_ = 1 if individual *i* is heterozygous for variant *j* (see Methods).

Using linear ARGs computed from UK Biobank (N=200k), we performed a common variant GWAS (MAF >0.01) over 89 traits and 42 covariates (Supplementary Table 4) (chromosomes 1-22, M=9,886,035 variants). We benchmarked our implementation against PLINK 2.0^35,36^, a highly optimized method, on a compute node with 256 GB of memory and 32 cores (both *kodama-gwas* and PLINK 2.0 are parallelized). *kodama-gwas* ran in 100 seconds, 4,700 times faster than PLINK 2.0 (Figure 4a). Its z-scores were highly concordant with PLINK 2.0, with a correlation of >0.9987 across traits with up to 75% missingness (Figure 4b and Supplementary Figure 1c). Effect sizes and standard errors were also concordant (Supplementary Figure 1a). Importantly, *kodama-gwas* was 110 times faster and also was better powered compared to the PLINK 2.0 qt-residualize, the faster configuration of PLINK 2.0, which ran in 11,000 seconds. PLINK 2.0 qt-residualize has smaller effective sample size in proportion to the genotypic variance explained by the covariates, resulting in larger standard errors and smaller chi-squared statistics (Supplementary Figure 1b).

**Figure 4:**
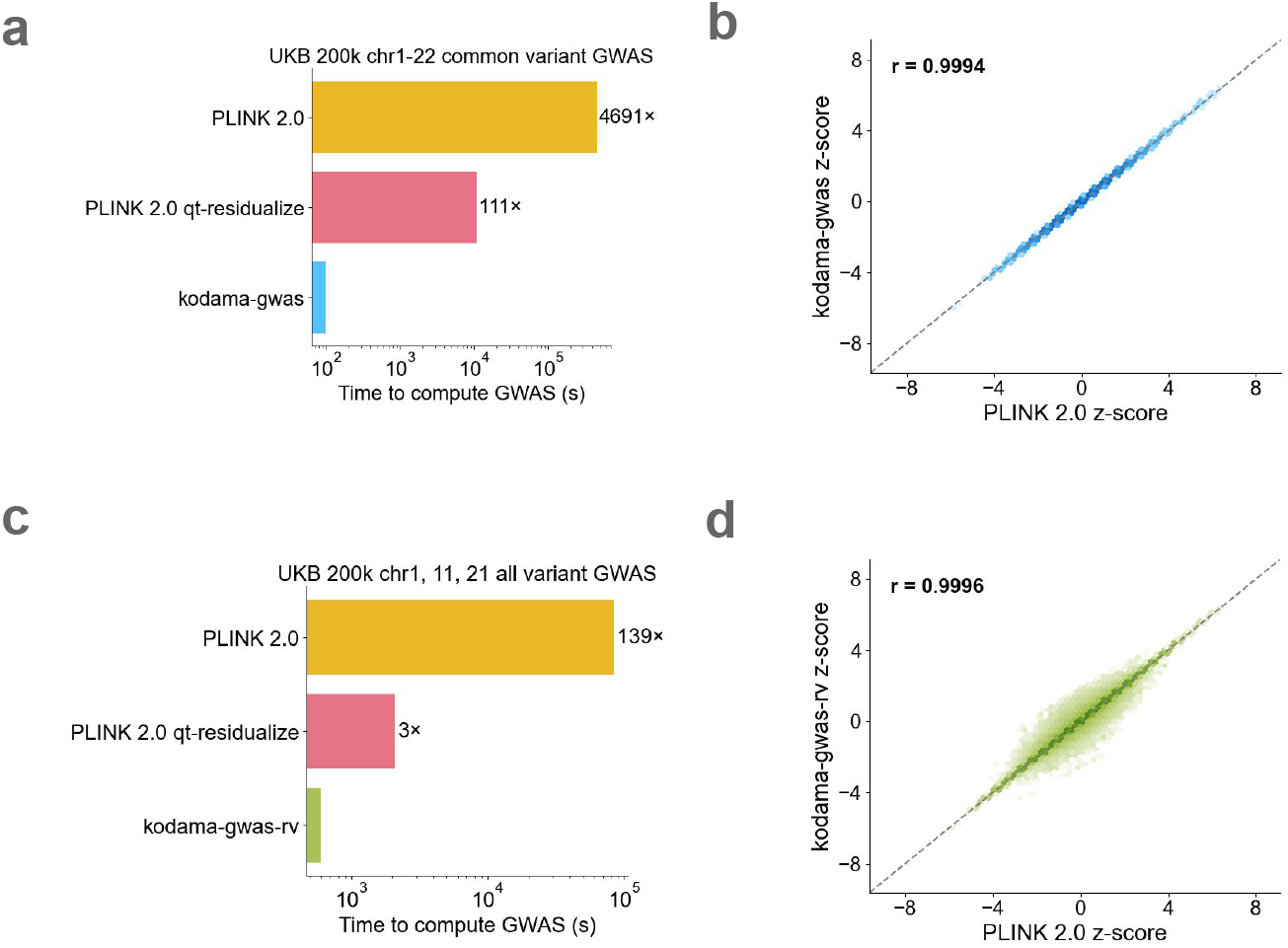
GWAS benchmark. **a, c** GWAS runtime comparison between the *kodama-gwas* and *kodama-gwas-rv* versus PLINK 2.0 on the UKB 200k common variants (chromosomes 1-22) and all variants (chromosomes 1, 11, 21) across 89 traits and 42 covariates. **b, d** z-score concordance between *kodama-gwas* and *kodama-gwas-rv* versus PLINK 2.0 for bilirubin (0.18 missingness) on chromosome 11. For numerical results, see Supplementary Tables 10 and 11.

We additionally performed an all variant GWAS using *kodama-gwas* and found that its results were discordant for some rare variants. This motivated us to use a second configuration of our method called *kodama-gwas-rv*, which unlike *kodama-gwas*, does not assume HWE and assumes that phenotypes are missing at random with respect to covariates but not with respect to genotypes by computing phenotype-specific allele counts (Methods). Although *kodama-gwas-rv* is slower than *kodama-gwas*, requiring 600 seconds to perform an all variant GWAS on chromosomes 1, 11, 21, it produces much more concordant results with PLINK 2.0 with a z-score correlation of >0.9984 across traits with up to 75% missingness on chromosome 11 (Supplementary Figure 2c). This represents a 140-fold speedup compared to PLINK 2.0 and a 3-fold speedup compared to PLINK 2.0 qt-residualize (Figure 4c).

GRG also implements an association scan, but we did not benchmark this method because it does not have the option to include covariates (we did benchmark the GRG matrix-vector operation; see below). Including covariates (especially principal components) is essential to avoid false positives in practice, and it greatly increases computational costs unless approximations are used.

### Genotype matrix operations

The association testing benchmark includes overhead from various sources, so we evaluated the performance of our matrix multiplication operations on their own, particularly in comparison with GRG. We also benchmarked a generic sparse matrix multiplication routine implemented in SciPy^37^ by loading in the data from .npz file format and then using the resulting CSC matrix to perform sparse matrix multiplication. Using UK Biobank data (N=200k) from chromosomes 1, 11, and 21, we benchmarked the time it took to load the data structure into memory and the time it took to perform left and right matrix multiplication. Compared to SciPy, the linear ARG loads into memory 32 times faster, and both matrix multiplications are approximately 40 times faster (Figure 5). Compared to GRG, loading the linear ARG is 2.5 times slower, but it performs matrix multiplication about 3 times faster (Figure 5). When multiple matrix multiplications are performed, which is common in practice, matrix multiplication runtime is larger than loading time, and the linear ARG is faster than GRG (Supplementary Figure 3a). Similarly, the linear ARG was 37 times faster than the PLINK 2.0 –score command and produced identical results (Supplementary Figure 3b).

**Figure 5:**
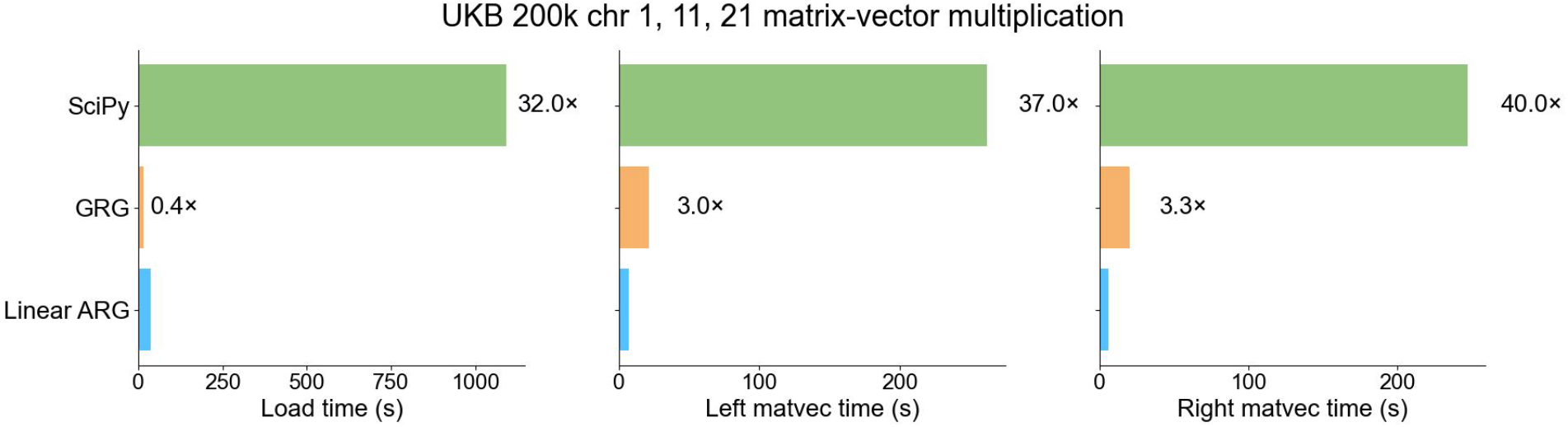
Matrix-vector multiplication benchmark. Matrix-vector CPU time comparison between the linear ARG, SciPy, and GRG on the UKB 200k across chromosomes 1, 11, 21. Load time is the time it takes to load each data structure into memory. Matrix-vector time is the time it takes to perform the matrix multiplication with a single vector. For numerical results, see Supplementary Table 12.

## Discussion

In genetic datasets, the amount of nonredundant information can be much smaller than the amount of raw data, because DNA sequences evolve slowly over time^8^. In comparative genomics, compressed data structures are widely used to search for sequence homology^38^. In human genetics, the Positional Burrows-Wheeler Transform (PBWT)^10^ has been a particularly successful compression method for applications like imputation and phasing^39–41^. ARG-based compression of simulated data has made it feasible to simulate realistic demographies at a massive scale^30,31^, and ARG inference has also become an important approach to demographic inference. In statistical genetics, inferred ARGs^21^ and the ARG-like GRG^9^ have begun to show promise but are not yet used widely. We introduced a genealogical compression method, *kodama*, that is designed for statistical applications like association scans. We showed *that kodama* achieves state-of-the-art compression ratios and reduces computational costs significantly in genetic association studies.

Our compressed data structure is closely related to the ancestral recombination graph (ARG), but with two key differences. First, the intermediate ‘brick graph’ data structure effectively pre-computes boundaries between segments of haplotypes that are inherited by different samples, creating a different node for each such segment. In the ARG, one haplotype can have different samples as descendants at different positions; in the brick graph, each node has a fixed set of descendants, and this property is convenient for matrix multiplication because it allows each node to store a single number. Second, the linearization step precomputes and subtracts redundant paths that arise due to a combination of recombination and coalescence. In principle, both of these differences increase the size of the linear ARG, such that an optimally compressed ARG should be smaller than an optimally compressed linear ARG; however, we believe that storing this additional information is beneficial when evaluating matrix products because otherwise, it needs to be recomputed implicitly. Because the linear ARG is defined based on the ARG (Supplementary Note), it is possible to convert any ARG into a linear ARG. While computing an ARG from a linear ARG (in particular, from the brick graph) may be possible, it will likely be more challenging because haplotype boundaries must be restored. Additionally, the brick graph contains no information about coalescence times, which would need to be inferred separately using a method such as *tsdate*^42^.

We note our approach is also related to the genotype representation graph (GRG)^9^. A GRG can be viewed as a brick graph with an added constraint, which is the number of paths between two nodes is 0 or 1. This constraint bypasses the redundancy problem that we solved instead using linearization. A downside of the GRG approach is that when a recombination event occurs between two haplotypes with a recent common ancestor, each of their common ancestors are bifurcated into two different nodes, increasing the size of the data structure (Supplementary Note). With our approach, common ancestors are not bifurcated, possibly explaining the ∼2.5x difference in compression (Figure 2a,b).

Linear ARGs implement the matrix-vector operations *Xβ* and *X′y*, and in many applications, these matrix products are the only way in which genotype data must be used. Performing a GWAS requires computing *X′y*; computing polygenic scores requires computing *Xβ*; RHE^43^ requires both kinds of matrix product. The fine mapping method SuSiE^44^ uses both matrix products within an iterative scheme where the effects of single variants are alternately removed from a vector of phenotype residuals, re-computed, and added back; linear ARGs may be especially beneficial in such applications because their small size allows them to be held in computer memory. Several methods, including REML and mixed model association methods, require solving the mixed model equations. A common way to do so is to use conjugate gradient^45^, an iterative method for solving linear equations that requires the matrix multiplication operation but not the matrix itself; such methods are called matrix-free.

Despite these benefits, other use cases of genotype data do not necessarily benefit from our approach. In particular, our data structure does not support random access to individual entries of the genotype matrix, because these entries are not stored; instead, they must be computed, and doing so has *O*(*N*) time complexity for common variants (Supplementary Note). The same limitation applies to the LD matrix *X*′*X* and the GRM *XX′*: these can be used as linear operators, but evaluating their individual matrix entries is inefficient. Some pipelines utilize linkage disequilibrium (LD) pruning, for computational reasons or otherwise; our data structure would have much less benefit after LD pruning, because LD is the reason that compression is possible. Finally, our data structure does not support the best-matching-haplotype lookup operations for which the PBWT is typically used, which are needed for imputation and phasing.

Another limitation of *kodama* is that it requires phased data with hard calls. It is not applicable to imputed dosages, and it also does not have any way to store information about missing or uncertain calls; this does limit the datasets to which it is applicable, but most large biobanks (particularly All of Us and UK Biobank) are performing WGS and making phased data available. Finally, the upfront cost of compression is a limitation. Compressing the UK Biobank N=200k dataset cost approximately £1,700 (Supplementary Table 3). This cost is worthwhile because of the savings in downstream analyses, but it could become problematic in extremely large datasets in the future. As such, we are making our linear ARGs available via UK Biobank Showcase for UK Biobank approved researchers (see Data Availability).

## Methods

### Linear ARG data structure

The linear ARG data structure consists of a sparse adjacency matrix *A*, two vectors specifying which nodes in *A* are variant and sample nodes, one vector specifying which variants have had their alleles flipped compared with the original genotype matrix, and a number of optional fields.

The adjacency matrix *A* encodes the ancestral relationship between nodes in the linear ARG. For example, if *A*_*ij*_ = *c* where *c* ≠ 0, there is an edge from node *i* to node *j* with edge weight *c*. This matrix is stored in SciPy compressed sparse column (CSC) format, which groups together the parents of each node. The CSC representation was chosen over CSR to enable a genotype matrix multiplication algorithm which re-uses the same memory addresses for multiple nodes (Supplementary Note).

The variant and sample indices are stored as NumPy^46^ arrays that specify which nodes in *A* correspond to variants and samples. These indices follow the ordering of the original genotype matrix *X*, such that the *i*-*th* element in the variant or sample index array refers to the *i*-*th* variant or sample in *X*.

The allele-flip vector is a boolean NumPy array that indicates which variants have been inverted in the linear ARG from reference to alternative alleles. These variants are flipped back to match the original genotype matrix when performing calculations.

The adjacency matrix *A*, the variant and sample indices, and flip are sufficient to compute matrix products with *X*. However, additional fields can also be loaded into the linear ARG data structure including:

- *iids*: NumPy array containing the sample ID for each haplotype; for diploid samples, this array contains two identcal entries
- *variants*: Polars lazyframe that stores variant metadata including CHROM, POS, ID, REF, and ALT
- *nonunique_indices*: NumPy array with a nonunique index for each node, used during matrix multplicaton to reduce memory consumpton (Supplementary Note)

Our Python API exposes genotype matrix multiplication on this data structure directly.

### Inferring linear ARGs from biobank-scale data

#### Multi-step compress pipeline

Due to the size of biobank-scale datasets, we implemented the multi-step compress pipeline that infers linear ARGs on smaller segments of the data and later merges them together. This approach reduces compute costs by reducing the memory required for most steps.

First, a large partition size is chosen so that each chromosome is divided into roughly equal segments of that size. A smaller partition size is selected to further divide each large partition into smaller segments of size approximately the small partition size. Second, for each small partition, a recombined brick graph is inferred (Steps 1 and 2 in Figure 1d). Third, all recombined brick graphs within each large partition are merged by first combining the graphs so that sample nodes are shared across small partitions, while any additional nodes are added as new nodes. Fourth, additional recombinations in the merged graph are found and the resulting graph is linearized (Steps 2 and 3 in Figure 1d). The large partitions are not further merged for memory efficiency and to allow for parallelization.

Partition sizes should be chosen carefully as larger partitions will result in higher memory usage, but also improve the compression ratio for two reasons: (1) the brick graph property may be able to exclude additional edges and (2) more recombinations between internal nodes may be found. The choice in partition size will depend on use case and memory constraints. Our choice of partition sizes for UK Biobank and All of Us were determined by identifying the smallest partition sizes that did not significantly hurt compression (See Datasets analyzed for more details).

### Verifying lossless compression

To check whether or not the inferred linear ARGs were lossless, we computed the allele counts using the genotype matrix and the linear ARG by left multiplying by the ones vector and confirmed that the allele counts were the same.

### Datasets analyzed

#### 1000 Genomes

We downloaded phased VCFs for 1000 Genomes (see Data Availability) and inferred linear ARGs using the single-step *kodama* compress command to each chromosome, on a laptop computer.

#### UK Biobank 200k

We inferred linear ARGs for chromosomes 1 to 22 by applying the multi-step compress pipeline using a large partition size of 20 Mb and a small partition size of 1 Mb. We ran this twice: once without any variant filtering, and once with a minor allele frequency (MAF) filter of >0.01.

For both the all variant and MAF filtered linear ARG, we inferred two additional linear ARGs with individual nodes needed for the no-HWE configuration. The total cost to infer the linear ARGs was £1,700 and £1,900 for the no filter and MAF filtered linear ARGs respectively (Supplementary Table 3). Although the no filter inference run took both more time and more memory, we were able to run these jobs on low priority while the MAF filtered inference runs were run as high priority jobs, explaining the difference in cost.

#### UK Biobank 500k

We applied our inference pipeline to the chromosome 21 of the 500k dataset. Since the DRAGEN VCF files contain multiallelic sites^2^, we first used “bcftools norm”^47^ to deduplicate the variants and then applied the multi-step compress pipeline (steps 1-3) using a large partition size of 20Mb and a small partition size of 0.5 Mb. All jobs were run on high priority and the total cost of inference was £100 (Supplementary Table 3).

#### All of Us

We also applied our inference pipeline to chromosome 21 of the AoU sr-WGS phased dataset (version 8, N=414,830; AC >= 2). The original sr-WGS phased vcf files contained multiallelic sites, however we chose not to deduplicate using bcftools, due to large initial write times, and implemented a “--remove-multiallelics” flag that filters these sites during parsing. We applied our multi-step compress pipeline using a large partition size of 20Mb and a small partition size of 1Mb. All jobs were run on high priority. The total cost to infer the linear ARG was no more than $141.88 (identifying exact costs is challenging on the AoU platform, hence an upper bound).

### GRG and XSI

To benchmark the linear ARG against other compression algorithms, we ran GRG and XSI on the same VCF files used to generate the linear ARG for chromosomes 1, 11, and 21. GRG was run using the recommended parameters corresponding to UK Biobank (3 trees and 50 parts for chromosomes 1, 4 trees and 240 parts for chromosome 11, 4 trees and 70 parts for chromosome 21) using version 2.0 while XSI was run with default parameters. The size of the .grg and .xsi files were used in the disk size comparison in Figure 2b.

### Testing optimality and robustness of the linear ARG with *msprime* simulations

We tested the optimality and the robustness of the linear ARG by simulating ARGs using *msprime*^30,31^. We simulated 100 ARGs under the OutOfAfrica_3G09 model^32^ in stdpopsim^33,34^ over the region chr21:21,990,355-23,324,211 with 10,000 and 100,000 samples under the infinite-sites assumption.

To test the optimality of the linear ARG, we extracted the genotype matrix from each simulated ARG and used it to infer a linear ARG. We compared the number of non-zeros in the linear ARG adjacency matrix to the number of edges in the simulated ARGs (ts.num_edges).

To test the robustness of the linear ARG compression ratio to mutations and errors, we simulated the following on the 10,000 sample genotype matrices extracted from the *msprime* simulations:

1. back mutation: uniformly select a variant and in the marginal tree of that variant, uniformly select a descendant edge such that all descendants of that edge no longer have that mutation
2. recurrent mutation: uniformly select a variant and in the marginal tree of that variant, uniformly select a non-descendant, non-ancestral edge such that all descendants of that edge have that mutation
3. genotyping error: uniformly select an individual and variant and flip the value in the genotype matrix from 0 to 1 or 1 to 0
4. out-of-order error: randomly select a pair of variants and switch their genomic positions

We simulated 0 to 50,000 events on each genotype matrix. Then we inferred the linear ARG from these mutated genotype matrices and quantified the brick graph compression ratio as the number of non-zeros in the simulated genotype matrix over the number of non-zeros in the brick graph and the linear ARG compression ratio as the number of non-zeros in the simulated genotype matrix over the number of non-zeros in the linear ARG.

### GWAS using the linear ARG

We implemented in *kodama-gwas* an association scan that accesses the genotype matrix only via matrix-vector products. Let *X* be the 2*n* × *m* phased genotype matrix, and let *y* be the 2*n* + 1 vector of phenotypes, with identical entries for the two haplotypes of each individual. Let *C* be the 2*n* × *k* matrix of covariates, which we assume has rank *k*. One of the covariates is the all-ones vector, corresponding to the intercept of the regression. First, we regress *C* out from *y*:

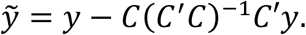

This calculation is fast because it has no dependence on the number of variants. Second, we compute the variance explained by *C* in each column of *X*:

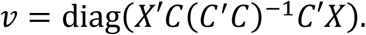

This calculation is performed using a single matrix-matrix product operation *z* = *X′C*, which for a single-trait GWAS is the most expensive step of the procedure. The diagonal of the matrix outer product is calculated using the identity:

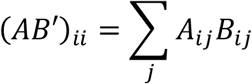

where *A* = *X′C* and *B′* = (*C′C*)^−1^ *X′C* (the pseudoinverse is used to handle rank-deficient *C*).

Third, we compute the product 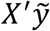 and compute the per-allele effect size of variant *j* as:

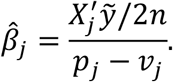

*p* is the allele frequency vector, which is the column of *X′C* corresponding to the all-ones covariate. When there are no covariates except for the all-ones covariate, then 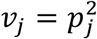, and the denominator is 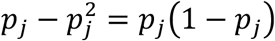. This formula assumes Hardy-Weinberg equilibrium (see below).

Fourth, we compute the standard error of 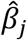, which is the square root of:

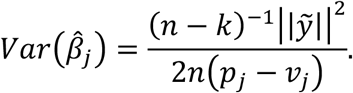

### Hardy-Weinberg equilibrium

The formulas in the previous section assume Hardy-Weinberg equilibrium conditional on covariates; this section identifies that assumption and shows how it can be avoided. Let *x*_1_, *x*_2_ ∈ {0,1}^n^ denote the paternal and maternal genotype vectors for some variant. Let *z*_1_, *z*_2_ ∈ ℝ^n^ contain the expected values of *x*_1_ and *x*_2_ conditional on covariates (i.e., the projections of *x* onto the span of *C*). Let 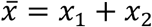 and let 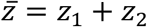. These should have the following properties:

1. *z*_1_ = *z*_2_, because covariates are identical for both haplotypes of the same individual
2. 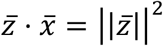, because the residuals 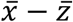 should be uncorrelated with the predictions 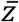

The GWAS effect size for this variant has the following denominator:

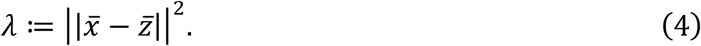

(The numerator is 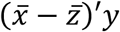, or equivalently 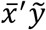). Using properties (1-2) above:

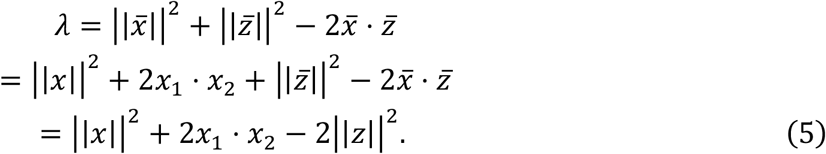

There are three terms:

1. ||*x*||^2^ is the allele count, 2*np*, because the genotypes are binary and 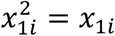 .
2. 2*x*_1_ ⋅ *x*_2_ is twice the number of homozygous carriers. The factor of two comes from the fact that for a homozygous sample *i*, with *x*_1*i*_ = *x*_2*i*_ = 1, the contribution to 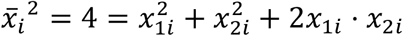.
3. ||z||^2^ is 2*nv* from the previous section. Because *z*_1_ = *z*_2_, it follows that 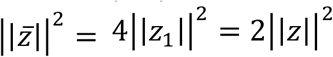.

Conditional Hardy-Weinberg equilibrium is the assumption that:

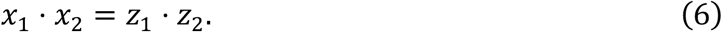

For example, if *z*_1_ = *z*_2_ = *p*, the allele frequency, then the fraction of homozygotes should equal *p*^2^. From (6) it follows that:

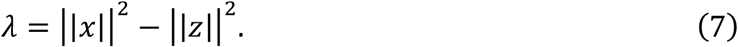

The GWAS effect size estimate is:

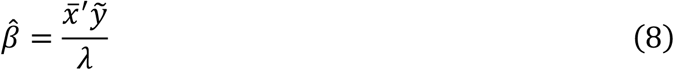

and the GWAS effect size variance is:

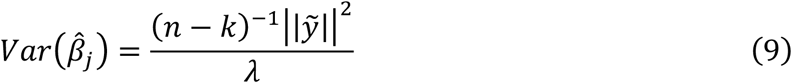

Substituting (6) for *λ* recovers equations (7)-(8).

We also implemented a configuration of *kodama-gwas* which avoids the Hardy-Weinberg assumption. This configuration is recommended for rare variants, which occasionally deviate from HWE by chance. This configuration requires computing the number of homozygotes, *x*_1_ ⋅ *x*_2_; then, it uses equation (5) instead of (7) for *λ*. Computing the number of homozygotes requires modifying the linear ARG data structure to include extra ‘individual nodes’, and then performing a matrix-vector product using the ordinary procedure; see Supplementary Note.

### Multiple phenotypes

Evaluating equations (8) and (9) requires the following components:

1. The residual matrix after projecting out covariates, 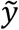
2. The allele count for each variant, ||*x*||^2^
3. The variance explained by covariates for each variant, ||z||^2^
4. If HWE is not assumed, the number of homozygotes, *x*_1_ ⋅ *x*_2_.

In principle, only item (1) is trait specific, and items (2-4) can be computed once and then re-used across traits. Doing this is beneficial because it is expensive to compute the matrix product *X′C*, which is used to compute ||z||^2^. However, a practical consideration is that phenotype values can be missing, and missingness may differ across traits; if *X′C* must be recomputed for each phenotype, it increases runtime from *O*(traits + covariates) to *O*(traits + covariates). This problem is specific to the denominator of the regression coefficients; the numerator can be computed by regressing phenotypes out from *Y*, imputing zeros for missing values, and calculating *X′Y*. The `--qt-residualize` configuration of PLINK 2.0 uses this approach for the numerator and ignores covariates when computing the denominator of the regression, but this approach overestimates the denominator (and underestimates effect sizes). REGENIE, which also batches computation across traits, does not use such an approximation; instead, it computes genotype residuals directly, possibly explaining the much higher runtime of REGENIE step 2 compared with PLINK 2.0 qt-residualize.

*kodama-gwas* implements two different ways of handling missingness which are both improvements over the qt-residualize approach. In its default, fastest configuration, it computes the denominator, *λ*, just once using the set of individuals who have a non-missing value for any phenotype. It scales this value by the fraction of non-missing individuals for each phenotype, effectively assuming that phenotypes are missing at random (MAR) with respect to both phenotypes and genotypes. If *l, k* are the number of phenotypes and covariates, then this approach requires *l* + *k* matrix-vector operations (*l* of them for the numerator, *k* of them for the denominator). It works well for common variants, but for rare variants, the MAR assumption will often fail for individual alleles due to chance; for example, if 10% of individuals are missing for some trait, then for a variant with allele count equal to two in the full sample, the allele count for that variant among nonmissing samples will be one or zero approximately 20% of the time. Using an allele frequency of 2/2*n* versus 1/(2*n* ∗ 0.9) is problematic.

For this reason, we provide a more accurate configuration, `recompute-AC`, which computes phenotype-specific allele counts. This configuration assumes that phenotypes are MAR with respect to covariates, but not with respect to genotypes. It requires 2*l* + *k* matrix-vector products, as for each phenotype we must compute both *X′y* and *X′*1_*y*_, where 1_*y*_ is an indicator vector for non-missing phenotypes. For some phenotype and for some variant, let *n* and *n′* be the whole-sample sample size and the non-missing sample size, let *p* and *p*′ be the whole-sample allele frequency and the non-missing allele frequency, and let *v* be the whole-sample estimate for the genotypic variance explained by covariates. Instead of computing 2*n*(*p* − *v*) for *λ*, we compute:

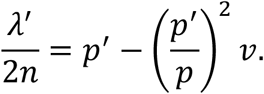

Notice that if there are no additional covariates besides the all-ones vector, then the variance explained is *v* = *p*^2^ (such that *p* − *v* = *p*(1 − *p*)), and the variance explained among non-missing samples is *v*^!^ = (*p*′)^2^, so this formula correctly gives us

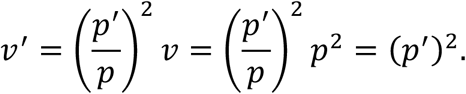

The `recompute-AC` configuration can be combined with the correction for Hardy-Weinberg disequilibrium. Let *q* be the whole-sample fraction of homozygotes. When these are combined, the formula approximating *λ* is:

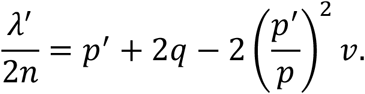

This formula is expected to be accurate for common variants by the law of large numbers, and it is expected to be accurate for rare variants because for rare variants, *q* ≪ *p*.

We recommend using the default (fastest) configuration of *kodama-gwas* when analyzing variants with minor allele count greater than 100 only and using both the `recompute-AC` and `no-hwe` options when analyzing rare variants or a combination of both rare and common variants.

### Benchmarking

#### Genome-wide association

##### Selection and preprocessing of traits and covariates

We selected 89 quantitative traits, prioritizing molecular traits, and selecting traits with less than 80% missingness (Supplementary Table 4). We included 42 covariates including age, sex, and the first 40 genetic PCs (Supplementary Table 4). The quantitative traits were averaged across instances when more than one measurement was available per individual and then quantile normalized. The covariates were mean centered and standardized. These traits and covariates were used for all GWAS runs.

##### Variant subsets

We benchmarked GWAS on the UK Biobank 200k dataset for chromosomes 1-22 on common variants (MAF > 0.01) and chromosomes 1, 11, and 21 for all variants.

##### PLINK 2.0

We performed GWAS using PLINK 2.0 with the *glm* function, PGEN files as input, the --omit-ref flag, and conducted benchmarks both with and without the --qt-residualize flag. We ran both of these configurations for the common and all variant GWAS.

##### kodama

We performed GWAS using *kodama* with the ‘*kodama assoc’* command. For the common variant GWAS, we ran *kodama-gwas* (default configuration) on the common variant linear ARG. For the all variant GWAS, we ran *kodama-gwas-rv* (--recompute-ac and --no-hwe flags) on the all variant linear ARG with individual nodes (Supplementary Note).

##### Computational resources

All GWAS runs were performed on a mem3_ssd1_v2_x32 node.

#### Matrix multiplication and PRS

We benchmarked the time it took for the linear ARG to perform right and left matrix multiplication on randomly generated matrices with dimensions 1 to 100 against SciPy sparse, GRG, and XSI across chromosomes 1, 11, 21. For the SciPy sparse, GRG, and the linear ARG, we measured the composite time it took to read in the data structures into memory and perform the matrix multiplication. For SciPy sparse, we read in the sparse matrix into memory from CSC format and performed matrix multiplication with the “@” symbol. For GRG, we loaded our inferred GRGs into memory using pygrgl.load_immutable_grg and then used pygrgl.matmul to perform matrix-matrix multiplication. For the linear ARG, we loaded the linear ARGs without variant metadata and performed the matrix multiplication using our graph traversal. We repeated this analysis for both right and left matrix multiplication. To benchmark XSI, we measured the time it took to run the dot_prod command which computes a left multiplication with a single vector. We quantified the time it took to perform the matrix multiplication as the sum of this time and the number of dimensions of the matrix as XSI does not support matrix multiplication. All benchmarks were performed on the same node: mem3_ssd1_v2_x16.

We implemented a PRS using the linear ARG, which has near-identical inputs (scores inputted as a .parquet instead of a .csv) and outputs to the PLINK 2.0 score function. We benchmarked our implementation with PLINK 2.0 on common variants across UKB 200k chromosomes 1-22 using randomly generated weights for a single trait. In our implementation, we scored all variants in a single run, while for PLINK 2.0, we scored each chromosome at a time and confirmed the sum of the scores across chromosomes was identical to the output of the linear ARG. Runtimes include the time it takes to load the variant metadata, perform variant filtering, and save the PRS scores. Both methods were run using a mem3_ssd1_v2_x32 node.

## Data availability

The 1000 Genomes phased vcf data is available at https://ftp.1000genomes.ebi.ac.uk/vol1/ftp/data_collections/1000G_2504_high_coverage/working/20220422_3202_phased_SNV_INDEL_SV/. UK Biobank data are available at http://www.ukbiobank.ac.uk. All of Us data are available to authorized users on the All of Us Researcher Workbench. The 1000 Genomes linear ARGs are publicly available for download at https://zenodo.org/records/18893386. UKB linear ARGs will be made available via the UKB-RAP Showcase for approved researchers.

## Code availability

*Kodama* is available at https://github.com/quattro/linear-dag/tree/main. Scripts used to perform the analyses and reproduce the figures are available at https://github.com/amberzshen/linear-arg-scripts. Scripts to perform the All of Us analysis are available at https://github.com/xinranwang111/kodama_aou/tree/main.

## Acknowledgements

We thank P. Palamara, A. Martin, S. Raychaudhuri, and H. Finucane for helpful discussions. This research was conducted using the UKB resource under application no. 86805. N.M. acknowledges funding from P01CA196569 and R01HG012133. L.J.O. acknowledges funding from the NIH NIGMS R35 GM155278.

