## Supplementary Note for "Highly efficient genotype compression leveraging genealogical relatedness"

The Supplementary Note is organized as follows. Section 1 defines the linear ancestral recombination graph and shows that any ARG corresponds to some linear ARG. Section 2 defines a graph, the ‘empirical brick graph’, which is computed in the first step of *kodama*. Sections 3-5 describe the ‘brick graph algorithm’, which computes the empirical brick graph; Section 3 justifies the approach, Section 4 describes the key algorithms, and Section 5 describes a post-processing step. Section 6 describes the second step of *kodama*, which involves adding “recombination nodes” to the empirical brick graph. Section 7 describes the third step of *kodama,* which is to “linearize” a brick graph by adding extra edges and possibly-negative edge weights. Section 8 describes how we perform genotype matrix operations on the linear ARG using a memory efficient approach. Section 9 describes a way to modify the linear ARG that allows us to compute the number of heterozygotes for each variant across any subset of individuals.

### 1. Definition of the linear ancestral recombination graph

An ancestral recombination graph (ARG) can be represented as a directed acyclic graph $G=\left( H,E \right)$ comprising a set of haplotypes as vertices and a set of edges that indicate local parentage. An edge between haplotypes $h_{1}$ and $h_{2}$ indicates that haplotype $h_{2}$ is a child of $h_{1}$. Edges are assigned genomic intervals via a function $\mathrm{int}:E\to\mathbb{N}^{2}$; an edge $e=(h_{1},h_{2})$ with $\mathrm{int}\left( e \right)=\left( a,b \right)$ indicates that $h_{1}$ is the parent of $h_{2}$ at positions $a,a+1, \ldots,b-1$. Every haplotype has at most one parent at any position: if $e_{1}=\left( h_{1},h_{3} \right)$ and $e_{2}=\left( h_{2},h_{3} \right)\in E$, and $h_{1}\neq h_{2}$, then $\mathrm{int}\left( e_{1} \right)\cap\mathrm{int}\left( e_{2} \right)=\emptyset$. Thus, the ARG can be regarded as a sequence of *marginal trees* (properly, forests) along the genome. This definition excludes coalescence times.

**Definition 1.** The **marginal tree** of an ARG $G=\left( H,E,\mathrm{int} \right)$ at position $x\mathbb{\in N}$ is the ARG $G\left( x \right)=\left( H,E\left( x \right),\mathrm{in}t_{x} \right)$ of $G$, where $E\left( x \right)=\{e\in E:x\in\mathrm{int}\left( e \right)\},$ and $\mathrm{int}_{x}: E\left( x \right)\to\left\{ x \right\}$ is constant.

Mutations can be placed upon an ARG. Under infinite sites, a unique derived allele at site $j$ arises at most once in the ARG as a mutation. The position of the site is $x=\mathrm{pos}(j)$ and the edge upon which the mutation occurs, if any, is $e=\mu\left( j \right)$, where $x\in\mathrm{int}\left( e \right)$. The carriers of the derived allele, $H\left( j \right)\subset H$, are the descendants of $e$ in the marginal tree $G\left( x \right)$. (The descendants of an edge $\left( u,v \right)$ are the haplotypes reachable from $v$, including $v$ itself). For a fixed set of sites $1,\ldots,m$, the genotype of a haplotype $h$ is a binary vector $X_{h}$ of length $m$, where $X_{h}\left( j \right)=1$ indicates that $h\in H\left( j \right)$. Sites are ordered such that if $i<j$, then $\mathrm{pos}\left( i \right)<\mathrm{pos}\left( j \right)$.

A challenge of working with the ARG is that different mutations upon the same edge can have different sets of carriers, depending upon their position. Nowbandegani & Wohns et al.^24^ introduced a modified ARG, which they called a bricked tree sequence and we call a *bricked ARG*, which is a directed multigraph with the same set of vertices (haplotypes) as the ARG but an expanded set of edges. In a multigraph, two vertices can have multiple edges. In the bricked ARG, an edge of the ARG is partitioned into multiple edges with the same parent and child haplotypes, but different genomic intervals. The partition is such that edges have the same descendant haplotypes at every position. These edges are called *bricks*.

**Definition 2.** The **bricked ARG** corresponding to an ARG $G=\left( H,E \right)$ is a multigraph $G^{'}=\left( H,B \right).$ For an edge $e=\left( u,v \right)\in E$, let $D_{e}\left( x \right)\subset H$ be the descendants of $v$ in $G\left( x \right)$. Let $P_{e}=\left[ x_{1},x_{2} \right), \left[ x_{2},x_{3} \right),\ldots$ be the minimal partition of $\mathrm{int}\left( e \right)$ such that $D_{e}\left( x \right)$ is constant on each element. $B$ is the union of edges $\left( e,\left[ x,y \right) \right)$ for every edge $e\in E$ and every interval $\left[ x,y \right)\in P_{e}$.

Just as the ARG can be sliced to extract the marginal tree at a position in the genome, a bricked ARG can be sliced to extract the ancestors of a given haplotype $h$.

**Definition 3.** In a bricked ARG $G^{'}=(H,B)$, the **brick diagram** $G_{h}^{'}=\left( H,B_{h} \right)$ of a haplotype $h$ is the bricked ARG whose edges $B_{h}$ are ancestors of $h$.

The brick diagram is defined specifically for a bricked ARG because in a general ARG, an edge could have $h$ as a descendant at one position but not a different position; this makes it ambiguous which edges ought to belong to $B_{h}$.

This simplest possible ARG is a *lineage*, with edges $\left( h_{1},h_{2} \right),\left( h_{2},h_{3} \right),\ldots,\left( h_{n-1},h_{n} \right)$. Brick diagrams have the following key property.

**Proposition 1 (*brick diagram property***). *A marginal tree of a brick diagram is a lineage.*

**Proof:** If two bricks $u,v\in B$ belong to the marginal tree at position $x$ of the brick diagram of haplotype $h$, then they are ancestors of $h$, and $x\in\mathrm{int}\left( u \right)\cap\mathrm{int}\left( v \right)$. Because $u$ and $v$ are ancestors of $h$ at every position in $\mathrm{int}(u)$ and $\mathrm{int}\left( v \right)$ respectively, they are both ancestors of $h$ in the marginal tree $G\left( x \right)$. Because $G\left( x \right)$ is a tree, either $u$ is an ancestor of $v$ or $v$ is an ancestor of $u$. This produces a total ordering for $B$.

The brick diagram property allows brick diagrams to be visualized in a 2-dimensional diagram resembling a “brick wall” (Figure). Nowbandegani & Wohns et al. previously defined the brick diagram for a pair of haplotypes instead of a single haplotype; our definition is equivalent to theirs in the case that one of the two is the root.

We define a graph whose nodes are not haplotypes but bricks:

**Definition 4**. For a bricked ARG $\left( H,B \right)$, the **brick graph** is the directed graph $\left( B,E \right)$ such that if $B$contains $b_{1}=\left( h_{1},h,{[x}_{1},y_{1}) \right)$ and $b_{2}=\left( h,h_{2},\left[ x_{2},y_{2} \right) \right)$, and either $x_{2}<y_{1}$ or $x_{1}<y_{2}$, then $\left( b_{1},b_{2} \right)\in E$.

In the brick graph, we say that a brick $u$ is an ancestor of a brick $v$ if there exists a path from $u$ to $v$, and $u\neq v$. We say that $u$ is an ancestor of a haplotype $x$ if there exists a path from $u$ to some brick $v$ whose child haplotype is $x$, possibly including $v=u$. The set of haplotypes which are descendants of a brick $u$ is denoted $H\left( u \right)$. By the brick diagram property, if $h\in H\left( u \right)$, then for any position $x\in\mathrm{int}\left( u \right)$ there exists a path $P$from $u$ to $h$ in the brick graph such that every $v\in P$ has $x\in\mathrm{int}\left( v \right).$

Bricks can be viewed as *pseudo-haplotypes* carrying certain alleles.

**Definition 5:** Let $G'=\left( H,B \right)$ be a bricked ARG, and let $\mu:\left[ m \right]\to B$ encode the position of mutations $1,\ldots,m$. The **pseudo-haplotype** of a brick $v\in B$ is the one-hot encoding $X_{v}\in\{0,1{\}}^{m}$ such that $X_{v}\left( j \right)=1$if, in the brick graph, $\mu\left( j \right)$ is an ancestor of $b$.

One aspect of this definition is artificial. A path in the brick graph may connect two bricks that exist at different intervals of the genome, and as a result, a brick $b$ may carry derived alleles that are not physically located within $\mathrm{int}\left( b \right)$. More generally, unlike haplotypes, pseudo-haplotypes do not correspond to segments of DNA that existed in the history of a population.

Despite this, pseudo-haplotypes are useful because they can be used to recover haplotypes.

**Proposition 2:** *Let* $X_{h}$ *be the one-hot encoding of mutations carried by haplotype* $h$*, and let* $B\left( h \right)\subset B$ *be the bricks whose child haplotype is* $h$*. Then:*

$$X_{h}=\sum_{v\in B(h)} X_{v}.$$

**Proof:** let $x=\mu\left( i \right)$ be the position of some mutation such that $X_{h}\left( i \right)=1$. $h$ is a descendant of $\mu\left( i \right)$. By the brick diagram property, the descendant bricks of $\mu\left( i \right)$ of which $h$ is a descendant form a lineage. Exactly one member of that lineage is a parent of $h$. Therefore, $\sum_{v\in B\left( h \right)} X_{v}\left( i \right)=1$.

Next, we show that pseudo-haplotypes are linear combinations of their ancestors – not only their parents, but also the common ancestors of their parents.

**Definition 6:** Let $G=(V,E)$ be a weighted directed acyclic graph, with edge weights $w:E\to R$ taking values in some ring (in particular, $\mathbb{Z}$ or $\mathcal{F}_{2}$). For $i,j\in V$, let $P\left( i,j \right)$ be the set of paths $\left( \left( i,k_{1} \right),\left( k_{1},k_{2} \right),\ldots,\left( k_{n},j \right) \right)$ from $i$ to $j$. The **path sum** from node $i$ to node $j$ is defined as the sum over paths of the product of its edge weights:

$$S\left( i,j \right)=\sum_{p\in P\left( i,j \right)} \prod_{e\in p} w\left( e \right).$$

The weighted DAG is called **1-summed** if for all $i,j\in V$, $S\left( i,j \right)=1$if $j\in\mathrm{reach}\left( i \right)$ and $S\left( i,j \right)=0$ otherwise.

**Lemma 1:** *Let* $A$ *be the adjacency matrix of a weighted DAG, and let* $S$ *be the matrix of path sums. Then:*

$$S=A+A^{2}+A^{3}+\cdots=A\left( I-A \right)^{-1}.$$

**Proof:** Each term in the series, $A^{k}$, is the path sum for paths of length $k$.

In particular, if the DAG is 1-summed, then entry $i,j$ of the matrix $A\left( I-A \right)^{-1}$ is an indicator for whether $j$ can be reached from $i$.

**Lemma 2**: *Given a DAG* $G=\left( V,E \right)$*, there exists a 1-summed DAG* $G^{+}=\left( V,E^{+} \right)$ *with nonzero edge weights* $w:E^{+}\to R$ *such that* $\mathrm{reach}\left( G^{+} \right)=\mathrm{reach}\left( G \right).$

**Proof**: This can be seen algebraically, but we construct $G^{+}$ directly from $G$ following our algorithm (see below). Label $V=\{1,\ldots,n\}$such that if $\left( i,j \right)\in E$, then $j<i$. Let $G^{\left( 0 \right)}=\left( V,E^{\left( 0 \right)} \right)=G$, with $w^{\left( 0 \right)}\left( e \right)=1$ for $e\in E^{\left( 0 \right)}$. Let $D\left( i \right)\subset V$ be the reachable set of $i$ in $G^{\left( i-1 \right)}$, and let $W\left( i,j \right)$ be the path sum from $i$ to $j$ in $G^{\left( i-1 \right)}.$ For $j\in D\left( i \right)$, let:

$$E^{\left( i \right)}=E^{\left( i-1 \right)}\cup\{\left( i,j \right): W\left( i,j \right)\neq1\},$$

and let $w^{\left( i \right)}:E^{\left( i \right)}\to R$ be:

$$w^{\left( i \right)}\left( e \right)=\left\{ \begin{matrix} w^{\left( i-1 \right)}\left( e \right)+1-W\left( i,j \right) & \mathrm{if} e\in E\left( i \right) \\ 1-W\left( i,j \right) & otherwise. \end{matrix} \right.$$

The path sum from $i$ to $j$ is the same in $G^{\left( i \right)}$ as it is in $G^{\left( n \right)}$, so $G^{(n)}$ is 1-summed.

**Definition 6:** Let $G=\left( B,E \right)$ be a brick graph. The **linear ancestral recombination graph** is the one-summed DAG $G^{+}$ such that $\mathrm{reach}\left( G^{+} \right)=\mathrm{reach}\left( G \right).$

**Theorem 1:** *Let* $G=(H,B)$ *be a brick graph with mutations* $[m]$*, and let* $M\in\{0,1{\}}^{|B|\times m}$ *be the matrix with ones in the entries* $\left( \mu\left( j \right),j \right)$ *and zeros elsewhere. Let* $X$ *be the genotype matrix, with one row* $X_{v}$ *for each brick* $v$*. Let* $A$ *be the adjacency matrix of* $G^{+}$*. Then:*

$$X=AX+M$$

*or equivalently,*

$$X=\left( I-A \right)^{-1}M.$$

**Proof:** By Lemma 1, the path sums in $G^{+}$ are:

$$S=A\left( I-A \right)^{-1}.$$

By Lemma 2, $S_{uv}$ is the indicator for whether $u$ is a descendent of $v$ in $G$. Moreover, $u$ has derived allele $j$ if $S_{u\mu\left( j \right)}=1$ or if $\mu(j)=u$. Therefore, $X$ can be written:

$$X=M+SM$$

$$=\left( I+A\left( I-A \right)^{-1} \right)M$$

$$=\left( I-A \right)^{-1}M.$$

### 2. The empirical brick graph

The first step of inferring the linear ARG is to compute the *empirical brick graph* of a phased genotype matrix. The emirical brick graph is a brick graph containing one brick for each variant. Let $G=\left( B,E \right)$ be a brick graph. Recall that the carriers of a mutation $j$, $H(j)$, are the haplotypes descended from $\mu\left( j \right)$ in $G$. In this section, the order of mutations in the genome is important; we assume that mutation indices follow the genome order, i.e., $i<j$ means that the position of $i$ is less than that of $j$.

**Lemma 3:** *Let* $i,j$ *be mutations such that* $\mu(i)$ *is an ancestor of* $\mu\left( k \right)$ *in the brick graph. Then* $H(k)\subset H(i)$*.*

**Proof.** There exists a path $P$ in the brick graph from $\mu(i)$ to $\mu(k)$. For each edge $\left( u,v \right)$ in $P$, $u$ and $v$ overlap at some position, and the descendant haplotypes of $v$ at that position are descendants of $u$ as well. Transitively, $H(k)\subset H(i)$.

**Theorem 2:** *Let* $i,j,k$*be mutations with* $i<j<k$*. Suppose that* $\mu(i)$ *is an ancestor of* $\mu\left( k \right)$ *in the brick graph, and that* $H(k)\cap H(j)\neq\emptyset$*. Then* $\mu\left( j \right)$ *is either an ancestor of* $\mu\left( k \right)$ *or a descendant of* $\mu\left( i \right)$*.*

**Proof.** Let $P\subset B$ be a path from $\mu(i)$ to $\mu(k)$, and let $h\in H\left( k \right)\cap H\left( j \right).$ Consider the brick diagram $G\left( h \right)$ (Figure). By Lemma 3, $G\left( h \right)$ contains all ancestors of $\mu\left( k \right)$, including the path $P.$ Some brick in $P$, $v$, has $\mathrm{int}\left( v \right)\cap int\left( \mu\left( j \right) \right)\neq\emptyset$. By the brick diagram property, if $v$ is not itself $\mu\left( j \right)$, then $\mu\left( j \right)$ is either an ancestor or a descendent of $v$. If an ancestor, then $\mu(j)$ is also an ancestor of $\mu\left( k \right)$, and $H(k)\subset H(j)$. If a descendent, then $\mu\left( i \right)$ is also a descendent of $\mu\left( i \right)$, and $H(j)\subset H(i)$.

Theorem 2 shows that the brick graph is locally tree-like, in the following sense. For any pair of nodes in a rooted tree, if they have any common descendant, then they are comparable (one of them is an ancestor of the other). The brick graph has a strictly weaker property, stating that if $i$ is comparable to $k$, then for any $j$ in $i,\ldots,k$ having a common descendant with both $i$ and $k$, $j$ is comparable with either $i$ or $k$.

Theorem 2 is useful for inference because it relates the unobserved structure of the brick graph to observed properties of the genotype matrix (namely, which variants are subsets of which other ones). We define the following estimator.

**Definition 7:** For a haploid genotype matrix $X\in\{{0,1\}}^{N\times M}$, the **empirical brick graph** is the transitive reduction $G=\left( \left[ M \right], E \right)$ of the graph $\left( \left[ M \right], \bar{E} \right)$ where $\left( i,k \right)\in\bar{E}$ if $i,j,k$satisfy Theorem 2 for all $j=i,i+1,\ldots,k$.

A naïve algorithm to compute the empirical brick graph would have runtime $O\left( NM^{3} \right)$: for every pair of variants $i,j$, verify the condition for every intervening variant $k$. In the following section, we describe an efficient algorithm to compute the empirical brick graph.

### 3. Justification for the brick graph algorithm

The brick graph algorithm computes the empirical brick graph in linear time, $O\left( NM \right)$. It is distantly related to the Positional Burrows-Wheeler Transform (PBWT), iterating over variants in their genomic order and maintaining a highly compressed summary of the information processed thus far. Whereas the PBWT maintains a permutation (ordering) of the samples, the brick graph algorithm maintains a tree with samples as leaves.

Let $T_{n}\subset2^{H}$ denote the “working tree” at position $n$, which is defined inductively as a function of $T_{n-1}$ and $H\left( n \right)$. Later, we will show that the elements of $T_{n}$ are nested subsets of each other, forming a tree. Let $s=s_{1},\ldots,s_{m}$ be a subsequence of the integers $\left[ n-1 \right]$. There exists a corresponding element of $T_{n-1}$, denoted $T_{n-1}\left( s \right)$. In the base case, $n=0,$ let $T_{0}\left( \emptyset\right)=H$. When $n>0$, we obtain a new subsequence of $[n]$ by taking a subsequence $s_{1},\ldots,s_{m}$ of $[n-1]$ and either appending $n$ or not. Let $X=T_{n-1}\left( s_{1},\ldots,s_{m} \right)$.

When we do append $n$:

$$T_{n}\left( s_{1},\ldots,s_{m},n \right)=X\cap H\left( n \right)$$

When we do not append $n$:

$$T_{n}\left( s_{1},\ldots,s_{m} \right)=\left\{ \begin{matrix} X & H\left( n \right)\subset X \\ X\cap H\left( n \right)^{c} & \mathrm{otherwise} \end{matrix} \right.$$

With this definition, $T_{n}\left( s \right)$ equals the intersection of $H\left( j \right)$ for $j\in s$, intersected with $H\left( j \right)^{c}$ for some but not all elements $j\leq n\notin s$.

**Theorem 3.1.** *A brick* $\mu\left( i \right)$ *is an ancestor of* $\mu(n+1)$*in the empirical brick graph if and only if there exists a subsequence* $s=i,s_{2},\ldots,s_{m}$ *of* $[n]$ *such that* $H\left( n+1 \right)\subset T_{n}\left( s \right)$*.*

We begin with the following.

**Lemma 4.** *Let* $s$ *be any subsequence of* $\left[ n \right]$ *such that for all* $i\in s$*,* $\mu\left( i \right)$ *is an ancestor of* $\mu\left( n+1 \right)$*. If* $T_{n}\left( s \right)\cap H\left( n+1 \right)$ *is nonempty, then* $H\left( n+1 \right)\subset T_{n}\left( s \right)$*.*

**Proof.** When $n=0$, $T_{0}=\{T_{0}\left( \emptyset\right)\}=\{H\}$. Suppose that Lemma 4 holds for all $k<n$; we will show that it holds for $k=n$. Let $s$ be any subsequence of $\left[ n \right]$ with $\mu\left( i \right)$ being an ancestor of $\mu\left( n+1 \right)$ for all $i\in s$.

Suppose that $j\notin s$, $s_{1}<j\leq n$. We will show that the effect of $j$ on $H\left( n+1 \right)$ does not cause a violation of the lemma because either:

A. $H\left( n+1 \right)\subset H\left( j \right)^{c}$, and the effect (intersecting with $H\left( j \right)^{c}$) does not cause a violation
B. $T_{j}\left( s\cap\left[ j \right] \right)=T_{j-1}\left( s\cap[j-1] \right)$, and there is no effect
C. $T_{j}\left( s\cap\left[ j \right] \right)\cap H\left( n+1 \right)=\emptyset$, and $T_{n}\left( s \right)\cap H\left( n+1 \right)$ is also empty.

By Theorem 2, there exist the following three possibilities:

1. $H\left( j \right)\cap H\left( n+1 \right)=\emptyset$. Then equivalently, $H\left( n+1 \right)\subset H\left( j \right)^{c}$ (case A).
2. $\mu\left( j \right)$ is an ancestor of $\mu\left( n+1 \right)$. If $H\left( j \right)$ is a subset of$T_{j-1}\left( s\cap\left[ j-1 \right] \right)$, then we have case B. Otherwise, $T_{j}\left( s\cap\left[ j \right] \right)\subset H\left( j \right)^{c}$, and because $H\left( n+1 \right)\subset H\left( j \right)$, $T_{j}\left( s\cap\left[ j \right] \right)$ is disjoint from $H\left( n+1 \right)$ (case C).
3. $\mu\left( j \right)$ is a descendant of $s_{k}$ for all $s_{k}\in s\cap\left[ j-1 \right]$. We may apply the inductive hypothesis to $s\cap[j-1]$ and $j$: if $T_{j-1}\left( s\cap[j-1] \right)\cap H\left( j \right)$ is empty, then $T_{j}\left( s\cap[j] \right)=T_{j-1}\left( s\cap[j-1] \right)\cap H\left( j \right)^{c}=T_{j-1}\left( s\cap\left[ j-1 \right] \right)$ (case B). If it is nonempty, then $H\left( j \right)\subset T_{j-1}\left( s\cap[j-1] \right)$ (also case B).

**Proof of Theorem 3.1**. Suppose that $\mu\left( i \right)$ is an ancestor of $\mu(n+1)$. Let $s$ be the subsequence of $i,\ldots,n$ that contains $j$ if $\mu\left( j \right)$ is an ancestor of $\mu\left( n+1 \right)$, such that $H\left( n+1 \right)\subset\bigcap_{j\in s} H\left( j \right)$. Suppose that some other $j<n\notin s$. If $H\left( j \right)$ is disjoint from $H\left( n+1 \right)$, then $H\left( n+1 \right)\subset H\left( j \right)^{c}$. Otherwise, by Theorem 2.1, $\mu\left( j \right)$ is a descendant of $s_{k}$ for all $s_{k}<j\in s$. By Lemma 4, either $H\left( j \right)$ is disjoint from $T_{j-1}\left( s\cap\left[ j-1 \right] \right)$, or $H\left( j \right)\subset T_{j-1}\left( s\cap\left[ j-1 \right] \right)$. In either case, $T_{j}\left( s\cap\left[ j \right] \right)=T_{j-1}\left( s\cap\left[ j-1 \right] \right)$. This suffices to show that $H\left( n+1 \right)\subset T_{n}\left( s \right)$.

Conversely, suppose that $H\left( n+1 \right)\subset T_{n}\left( s \right)$ for some $s$. Then $H\left( n+1 \right)\subset H\left( s_{k} \right)$ for all $s_{k}\in s$. For all $j\notin s$, $s_{1}<j\leq n$, either $H\left( j \right)\subset T_{j-1}\left( s\cap\left[ j-1 \right] \right)$, or $H\left( j \right)\cap H\left( n+1 \right)=\emptyset$. In the former case, suppose that Theorem 2.2 holds for all $i\leq n$, particularly including $j$. Applying it to $\mu\left( j \right)$ and the subsequence $s\cap\left[ j-1 \right]$, $\mu\left( s_{1} \right)$ must be an ancestor of $\mu\left( j \right)$. These three cases show that $s_{1}$ and $n+1$satisfy the condition of Theorem 2.1, and that ${\mu(s}_{1})$ is an ancestor of $\mu\left( n+1 \right)$.

**Theorem 3.2.** For any two elements of $T_{n}$, either they are disjoint, or one is a subset of the other.

**Proof.** The statement holds for $T_{0}$. Suppose that it holds for $T_{n-1}$. First, if $T_{n-1}(s)$, $T_{n-1}\left( s^{'} \right)$ are two elements of $T_{n}$ such that $T_{n-1}\left( s \right)\cap T_{n-1}\left( s^{'} \right)=\emptyset$, then $T_{n}\left( s \right)\cap T_{n}\left( s^{'} \right)=\emptyset$. The same is true if $n$ is appended to either $s$, $s'$, or both of them. Second, suppose that $T_{n-1}\left( s \right)\subset T_{n-1}\left( s^{'} \right).$ There are several cases to check:

1. If $n$ is appended to both subsequences, then $T_{n}\left( s,n \right)=T_{n-1}\left( s \right)\cap H\left( n \right)\subset T_{n-1}\left( s' \right)\cap H\left( n \right)=T_{n}\left( s^{'},n \right)$.
2. If $n$ is not appended to either subsequence, and $H\left( n \right)\subset T_{n}(s)$, then $H\left( n \right)\subset T_{n}\left( s \right)$ as well, and $T_{n}\left( s \right)=T_{n-1}\left( s \right)\subset T_{n-1}\left( s' \right)=T_{n}\left( s^{'} \right)$.
3. If $n$ is not appended to either subsequence, and $H\left( n \right)\subset T_{n-1}(s')$, then $T_{n}\left( s^{'} \right)=T_{n-1}\left( s^{'} \right)$ and $T_{n}\left( s^{'} \right)\subset T_{n-1}\left( s^{'} \right)\subset T_{n}\left( s^{'} \right)$.
4. If $n$ is not appended to either subsequence, and $H\left( n \right)$ is not a subset of $T_{n}\left( s' \right)$ or $T_{n}\left( s \right)$, then $T_{n}\left( s \right)=T_{n-1}\left( s \right)\cap H\left( n \right)^{c}\subset T_{n-1}\left( s^{'} \right)\cap H\left( n \right)^{c}=T_{n}\left( s^{'} \right)$.
5. If $n$ is appended to $s'$ but not $s$, and $H\left( n \right)\subset T_{n-1}\left( s \right)$, then $T_{n}\left( s^{'},n \right)=T_{n-1}\left( s^{'} \right)\cap H\left( n \right)\subset H\left( n \right)\subset T_{n-1}\left( s \right)=T_{n}\left( s \right)$.
6. If $n$ is appended to $s'$ but not $s$, and $H\left( n \right)$ is not a subset of $T_{n-1}\left( s \right)$, then $T_{n}\left( s^{'},n \right)\subset H\left( n \right)$ while $T_{n}\left( s \right)\subset H\left( n \right)^{c}$, so they are disjoint.
7. If $n$ is appended to $s$ but not $s'$, and $H\left( n \right)\subset T_{n-1}\left( s' \right)$, then $T_{n}\left( s,n \right)=T_{n-1}\left( s \right)\cap H\left( n \right)\subset T_{n-1}\left( s' \right)=T_{n}\left( s^{'},n \right)$.
8. If $n$ is appended to $s$ but not $s'$, and $H\left( n \right)$ is not a subset of $T_{n-1}\left( s \right)$, then $T_{n}\left( s,n \right)\subset H\left( n \right)$ while $T_{n}\left( s' \right)\subset H\left( n \right)^{c}$, so they are disjoint.

Every pair of elements in $T_{n}$ falls into one of these cases.

### 4. The brick graph algorithm

Theorem 3.2 shows that $T_{n}$ can be structured as a rooted tree, with $T_{n}\left( \emptyset\right)=H$ as the root, such that the descendants of a node are subsets of it. We take advantage of this fact in the brick graph algorithm. We begin by describing a simplified algorithm that demonstrates the core logic.

**Algorithm 1:** computes $T_{n+1}$ from $T_{n}$.

**Input:** a directed tree $T_{n}$ and a subset of its leaf nodes, $H\left( n+1 \right)$

Let $v_{1},\ldots,v_{m}$ be a sorted sequence of nodes in $T_{n}$, containing all ancestors of the leaves in $H\left( n+1 \right)$, such that if $v_{i}$ is a descendant of $v_{j}$ then $i<j$. Let $V=\left\{ v_{1},\ldots,v_{m} \right\}$ .

Let $T_{n+1}=T_{n}.$

For $i=1,\ldots,m$:

If all children of $v_{i}$ are in $V$ or $V^{c}$, continue

Create a new node $u$and add $u$ as a child of the unique parent of $v_{i}$ in $T_{n+1}$

For all children $w$ of $v_{i}$, if $w\notin V$, assign $u$ instead of $v_{i}$ as the parent of $w$ in $T_{n+1}$

**Return:** $T_{n+1}$

This procedure effectively takes the subtree of $T$ that is descended from the lowest common ancestor of $X$, splits it into two pieces (one a subset of $H\left( n+1 \right)$, one of its complement), and reattaches those two subtrees at the lowest common ancestor. It can be confirmed that if the descendants of a node $u$ in $T_{n}$ are equal to $T_{n}\left( s \right)$ for some sequence $s$, then $T_{n+1}$ contains nodes whose descendants are equal to both $T_{n+1}\left( s,n+1 \right)$ and $T_{n+1}(s)$.

Moreover, Algorithm 1 can be modified to identify all ancestors $i<n+1$ in the empirical brick graph. By Theorem 3.1, $i$ is an ancestor of $n+1$ if for some $s=i,s_{2},\ldots,s_{m}$ of $[n]$, $H\left( n+1 \right)\subset T_{n}\left( s \right)$. We must maintain a mapping from the node $u=T_{n}\left( s \right)$ to the variant $i$. In tree $T_{i}$, the element $T_{i}\left( i \right)$ contains $H\left( i \right)$; it corresponds to the lowest common ancestor $u$ of the leaves in $H\left( i \right)$, which form a clade in that tree. Thus, we label the lowest common ancestor of $H\left( i \right)$ with $i$. Subsequently, the clade may be bifurcated into smaller clades, and these retain the label $i$ (in addition to any new labels), until all nodes labeled $i$ are the singleton leaf nodes of $H\left( i \right)$. This procedure is described in Algorithm 2.

**Algorithm 2:** computes $T_{n+1}$ from $T_{n}$ and updates a node-to-variant mapping $S$.

**Input:** a directed tree $T_{n}$, a subset of its leaf nodes, $H\left( n+1 \right)$, and a mapping $S$ from the nodes of $T_{n}$ to $\{\emptyset\}\cup[n]$.

Let $v_{1},\ldots,v_{m}$ be a sorted sequence of nodes in $T_{n}$, containing all ancestors of the leaves in $H\left( n+1 \right)$, such that if $v_{i}$ is a descendant of $v_{j}$, then $i<j$. Let $V=\left\{ v_{1},\ldots,v_{m} \right\}$ .

Let $T_{n+1}=T_{n}.$

For $i=1,\ldots,m$:

If all children of $v_{i}$ are in $V$ or $V^{c}$, continue

Create a new node $u$and add $u$ as a child of the unique parent of $v_{i}$ in $T_{n+1}$

For all children $w$ of $v_{i}$, if $w\notin V$, assign $u$ instead of $v_{i}$ as the parent of $w$ in $T_{n+1}$

Copy $S\left( u \right)=S(v_{i})$

If $v_{i}$ is the lowest common ancestor of $H(n+1)$, append $n+1$ to $S\left( v_{i} \right)$

**Return:** $T_{n+1}$

After running Algorithm 2, we can identify preceding ancestors of node $n+1$ using Algorithm 3:

**Algorithm 3**: Computes the preceding ancestors of variant $n+1$.

**Input**: The mapping $S$, the tree $T_{n+1}$, and the LCA $u_{1}$ of $H(n+1)$

Let $u_{1},\ldots,u_{n}$ be the unique path from $u_{1}$ to the root of $T_{n+1}$

Let $A=\{\}$ be the preceding ancestors of variant $n+1$

For $i=1,\ldots,n$:

For all $j\in S\left( u_{i} \right)$, add $j$ to $A$

**Return:** $A$

By running Algorithms 2-3 on nodes $1,2,\ldots,N$ and then $N,N-1,\ldots,1$, we can compute all ancestors of all nodes. However, this procedure computes all ancestor-descendant pairs in the empirical brick graph, as opposed to parent-child pairs (i.e., it computes the transitive closure). One could do this and then take the transitive reduction, but this approach is prohibitively slow and memory-intensive on biobank-scale data. We bypass this issue by computing parent-child pairs directly, although as we will see, this leads to added complexity.

Specifically, we modify Algorithm 3 so that it directly computes the *forward reduction*, which is defined as the graph with an edge $\left( i,j \right)$ if:

- $i<j$
- $i$ is an ancestor of $j$
- There exists no third node $k$, $i<k<j$, such that $i$ is an ancestor of $k$ and $k$ is an ancestor of $j$.

Concisely, the forward reduction is the transitive reduction of the intersection of the partial ordering with the total ordering “$<"$.

**Algorithm 4**: updates the forward reduction, $G^{+}$.

**Input**: The mapping $S$, the tree $T_{n+1}$, and the LCA $u_{1}$ of $H(n+1)$

Let $u_{1},\ldots,u_{n}$ be the unique path from $u_{1}$ to the root of $T_{n+1}$

Let $A=\{\}$ be the preceding ancestors of variant $n+1$

Let $k_{\max}=-\infty$

For $j=1,\ldots,n:$

Let $k=\max\left( S\left( u_{j} \right) \right)$

If $k>k_{\max}$:

Add $\left( S\left( j \right), i \right)$ as an edge of $G^{+}$

Set $k_{\max}=k$

In the reverse direction, $k_{\max}$ is replaced by $k_{\min}$, it is initialized at $+\infty$, and the comparison is reversed.

This procedure works for the following reason. In $T_{n+1}$, suppose that $i\in S\left( u \right)$ and $j\in S\left( v \right)$, where $i<j$ and $u$ is an ancestor of $v$ in the tree. Then $i$ is a preceding ancestor of $j$. If $u_{1}$ is a mutual descendent of $u,v$, then in Algorithm 4, we encounter $v$ before $u$. We make an edge (potentially) with $j$ but not with $i$ because when reaching $i$, $k_{max}\geq j>i$.

The following algorithms combine the steps we have described thus far, in addition to a final step described below.

**Algorithm 5:** performs the forward pass.

**Input:** the genotype data, $H\left( 1 \right),\ldots,H\left( M \right)$

Let $T$ be a tree with $N+1$ nodes, one leaf node for each sample and a root node connected to every leaf

Let $S$ map each node $v\in T$ to one or zero variants; initially, $S\left( v \right)=\emptyset$

Let $G^{+}$ be the empty graph

For $i=1,\ldots,M:$

Let $v$ be the lowest common ancestor of $H(i)$ in $T$

Apply Algorithm 2 to update $T$ and $S$

Apply Algorithm 4 to $S, T$, and the LCA $v$ of $H\left( i \right)$, updating $G^{+}$

**Return**: $G^{+}, S, T$

**Algorithm 6:** the brick graph algorithm.

**Input:** the genotype data, $H\left( 1 \right),\ldots,H\left( M \right)$

Let $G^{+}, S, T$ be the output of the forward pass (Algorithm 5)

Label the samples in $H$ with integers $i>M$

For $i\in H$:

Let $v$ be the leaf node corresponding to sample $i$

Apply Algorithm 4 to $S, T$ and $v$, updating $G^{+}$

Let $G^{-}$ be the output of the backward pass (Algorithm 5 with variant order reversed)

Let $G$ be the output of Algorithm 7 (see below) applied to $G^{+}$ and $G^{-}$

**Return:** the empirical brick graph, $G$

In between the forward and the backward passes, sample nodes have been added to the graph, effectively extending the forward pass. These are equivalent to singleton mutations, and they can be skipped in the backward pass because they are never ancestors of other nodes. The variants that are carried by each sample are those which are ancestors of the sample in the brick graph.

### 5. Combining the forward and backward passes

The forward and backward passes of the brick graph algorithm compute the graphs $G^{+}$ and $G^{-}$, respectively. They are related to the empirical brick graph $G$ as follows. Let $P$ be the partial order encoded by $G$: $\left( i,j \right)\in P$ if there exists a path from $i$ to $j$ in $G$. We assume that there are no “ties”, i.e., variants corresponding to the same node of the brick graph (otherwise, $P$ is a *postorder* instead of a partial order; see below). Let

$$P^{+}="P\cap\text{>" }\text{≔}\{\left( i,j \right)\in P:i>j\},$$

and let $P^{-}=P\cap\text{<}.$ The graph computed by the forward pass, $G^{+}$, is the transitive reduction of $P^{+}$; $G^{-}$ is the transitive reduction of $P^{-}$.

Whereas $P=P^{+}\cup P^{-}$, it is false that $G=G^{+}\cup G^{-}$, because $G^{+}$ may contain $G$-transitive edges $\left( u,v \right)$ such that $G$, but not $G^{+}$, contains an alternative path from $u$ to $v$. However, the following shows that we can efficiently compute $G$ from $G^{+}$ and $G^{-}$.

**Proposition 3**. Suppose that $\left( u,v \right)$ is an edge in $G^{+}$ but not $G$. Then there exists a node $w$ such that either $\left( u,w \right)\in G^{+}$ and $\left( w,v \right)\in G^{-}$, or $\left( u,w \right)\in G^{-}$ and $\left( w,v \right)\in G^{+}$.

**Proof.** Let $u_{1}≽u_{2}$ denote that $\left( u_{1},u_{2} \right)\in P$. There exists a path $u,w_{1},\ldots,w_{n},v$ from $u$ to $v$ in $G$, with $n>0$. Every $w_{k}$ must be outside of the interval $[u,v]$ in the total ordering, because otherwise $\left( u,v \right)$ would not belong to $G^{+}$. In particular, for some $w_{i}$, either $w_{i}<u$ or $w_{i}>v$.

If $w_{i}<u$, let $W=\{x<u\in V:u≽x≽v\}$; $W$ is nonempty, as it contains $w_{i}.$ Let $w=\max\left( W \right)$. Then $u≽w$ and $u>w$, so $\left( u,w \right)\in P^{-}$. Moreover, there is no larger node $w^{'}$ such that $u≽w^{'}≽w$ and $u>w^{'}>w$, because this node would belong to $W$, but $w=\max\left( W \right)$. Therefore, $\left( u,w \right)\in G^{-}$. Moreover, $\left( w,v \right)\in G^{+}$, for the same reason.

If $w_{i}>v$, let $W=\{x>v\in V:u≽x≽v\}\neq\emptyset$, and let $w=\min\left( W \right)$; by the same argument, $\left( u,w \right)\in G^{+}$ and $\left( w,v \right)\in G^{-}$.

This result justifies the following algorithm.

**Algorithm 7:** Computes $G$ from $G^{+}$ and $G^{-}$.

**Input**: $G^{+}$ and $G^{-}$, the output of the forward and backward passes

Let $G$ be the graph with the same nodes $V$ as $G^{+}$, and no edges

For nodes $u\in V$:

Let $X^{+}$ be the children of $u$ in $G^{+}$

Let $S^{+}$ be the parents of any $x\in X^{+}$ in $G^{-}$

Let $X^{-}$ be the children of $u$ in $G^{-}$

Let $S^{-}$ be the parents of any $x\in X^{-}$ in $G^{+}$

Add $\left( u,v \right)$ to $G$ for all $v\in X^{+}\cup X^{-}$ if $v\notin S^{+}\cup S^{-}$

**Return:** $G$

Real data does include pairs $u,v$ with $\left( u,v \right)\in G^{+}$ and $\left( v,u \right)\in G^{-}$. These pairs correspond to variants with the exact same set of carriers. They are handled by finding all maximal sequences $u<v<\ldots<w<x$ that are connected via edges $\left( u,v \right),\ldots,\left( w,x \right)$ in $G^{+}$ and via $\left( x,w \right),\ldots,\left( v,u \right)$ in $G^{-}$. These sequences are combined into a single node before applying Algorithm 7; we describe this procedure informally.

If some pair of nodes $u,v$ has $\left( u,v \right)\in G^{+}$ and $\left( v,u \right)\in G^{-}$, then $v$ is the largest out-neighbor of $u$, and vice versa. We iterate through nodes $u$ in their forward genomic order and check whether the largest out-neighbor $v$ of $u$ in $G^{+}$ has $u$ as its smallest out-neighbor in $G^{-}$. If so, then we contract the edge between $u$ and $v$, adding some neighbors $w<u$ of $u$ as neighbors of $v$. Not all such neighbors should be added; some in-neighbors $w$ of $u$ in $G^{+}$ might have an alternative path to $v$ in $G^{+}$, of the form

$$w\to w^{'}\to\cdots\to v.$$

If such a path exists, then $w^{'}>u$. Because $u,v$ have the same ancestors in $G$, $\left( w^{'},u \right)$ must be an edge within $G^{-}$. Thus, when adding in-neighbors $w$ of $u$ to $v$ in $G^{+}$, we verify that the out-neighbors $w'$ of $w$ do not have an edge $\left( w^{'},u \right)\in G^{-}$. Likewise, in $G^{-}$, out-neighbors $w$ of $u$ are only added as out-neighbors of $v$ if the in-neighbors $w'$ of $w$ in $G^{-}$ do not have an edge $\left( u,w^{'} \right)\in G^{+}$.

In addition, contracting $\left( u,v \right)$ breaks paths $w\to u\to w^{'}$ in $G^{+}$ if $w^{'}<v$. (This path would still exist in $G^{+}\cup G^{-}$, but it is essential that the path exist in $G^{+}$ specifically). These paths are patched by adding edges between some in-neighbors $w$ of $u$ and out-neighbors $w^{'}<v$ in $G^{+}$. However, an alternative path may already exist in $G^{+}$. We search all descendants of $w$ in $G^{+}$ that are less than or equal to $w'$, excluding paths through $u$. If $w'$ is not found, then $\left( w,w^{'} \right)$ is added to $G^{+}$. We do the same for $G^{-}$.

After contracting an undirected edge, we keep vectors encoding node re-assignments, $v_{1}\sim v_{2}\sim\cdots\sim v_{k}.$ The node $v_{k}$ is assigned all of the variants that correspond to nodes $v_{1},\ldots,v_{k}$.

### 6. Finding recombination nodes

The empirical brick graph contains at most one node per variant (and sample). Missing from the graph are ancestral recombination events that do not coincide with any single mutation, and in their place, the graph contains inefficient *bicliques* connecting the recombinant parents with their children. A biclique comprises two sets of nodes, $U$ and $V$, such that $\left( u,v \right)\in E$ for all $u\in U$, $v\in V$. The number of edges in a biclique is $\left| U \right|\left| V \right|$, which can be large if a large number of children have several shared parents. Introducing a new recombination node, a biclique can be replaced by a “star” having $\left| U \right|+|V|$ edges (Main text Figure 1d).

In general, enumerating bicliques is challenging (for example, finding the largest biclique in a graph is NP-complete). However, recombinations only occur between haplotypes that are physically overlapping (or bricks that are physically adjacent), which allows us to restrict our search. Starting with the empirical brick graph, we begin by ordering the bricks $B$ according to the genomic position of their associated mutations. (The position of the samples can be anything). We define an *adjacent trio* as the tuple $\left( p_{1},p_{2},c \right)$ such that $e_{1}=\left( p_{1},c \right),e_{2}=\left( p_{2},c \right)\in E$, $p_{1}<p_{2}$, and if any $e_{3}=\left( p_{3},c \right)\in E$, then either $p_{3}\leq p_{1}$ or $p_{3}\geq p_{2}$. The number of adjacent trios is linear in the number of edges, as each edge can participate in at most two trios. We enumerate these trios, collecting them into bicliques involving two parents and any number of children. We iterate over bicliques from largest to smallest, replacing each with a “star”. This operation disturbs adjacent trios of the form $\left( p_{0},p_{1},c \right)$ or $\left( p_{2},p_{3},c \right)$; such trios are removed from their respective bicliques and assigned to a new biclique of the form $\left( p_{0},u,c \right)$ or $\left( u,p_{3},c \right)$. We maintain sorted linked lists of child nodes belonging to each biclique; this allows us to remove these trios from their respective bicliques in linear time (iterating once over the list).

This algorithm also requires that we track the order of each node’s parents. Each node in the graph has two pointers, to a linked list of in-edges and a linked list of out-edges. Each edge has six pointers: to its nodes, $u$ and $v$; to the next in-edge and the previous in-edge of the in-edge linked list of node $v$; and to the next out-edge and the previous out-edge of the out-edge linked list of node $u$. In-edge linked lists are maintained in the genomic order; for two variants $a,b$, if they occur at sites $x_{a}<x_{b}$ in the genome, then $a$ comes before $b$ in the linked list of in-edges for any shared child node. This way, trios of the form $\left( p_{1},p_{2},c \right)$ involve adjacent pairs of in-edges of $c$. Out-edge linked lists are in an arbitrary order.

The order of biclique elimination affects the graph that is produced. Our recombination-finding algorithm processes bicliques in order of their number of child nodes, such that if the child nodes of one biclique form a strict superset of another, then it will be processed before its subset biclique. This choice is not guaranteed to produce a maximally sparse graph in general, but it can be justified heuristically under a strong assumption, that identical haplotypes are always identical by descent (IBD) as opposed to by state (IBS).

Suppose that for every genomic interval, if two haplotypes $h_{1}$ and $h_{2}$ carry the same alleles within that interval, then their lowest common ancestor is also identical at every position in the interval. That is, identical haplotypes are IBD. Suppose that some haplotype $h_{1}$ carries the alleles $a,b,c$ at consecutive sites, another haplotype $h_{2}$ carries $a,b$, and $h_{3}$carries $b,c.$ Then $h_{1}$ and $h_{2}$ inherit $b$ from a common ancestor $h_{2}'$ which also has $a$; $h_{1},h_{3}$ inherit $b$ from a common ancestor $h_{3}'$ which also has $c$. If $h_{2}^{'}\neq h_{3}'$, then $h_{2}'$ is either an ancestor or a descendant of $h_{3}'$, because both are ancestors of $h_{1}$ at the site of allele $b$. Therefore, the carriers of $a,b$ are either a superset or a subset of the carriers of $b,c$. If they are strictly a superset, then they are factored first, producing the smallest possible graph:

$$h_{2}\leftarrow h_{2}^{'}\to h_{3}^{'}\to h_{1},h_{3}.$$

The no-IBS assumption is unlikely to hold in practice, such that our recombination-finding algorithm is almost certainly suboptimal.

### 7. Linearizing the brick graph

Although it is not the approach that we take, a brick graph can be turned into a linear ARG algebraically. Let $\tilde{X}$ be the square matrix such that $\tilde{X}_{ij}=1$ if node $i$ is reachable from node $j$ in the linear ARG (this has the genotype matrix $X$ as a submatrix). Then:

$$\begin{aligned} A=I-\tilde{X}^{-1}. \#\left( xy \right) \end{aligned}$$

In principle, it is straightforward to invert $\tilde{X}$ because it is triangular. In practice, on biobank-scale data, $\tilde{X}$ is too large to be stored in memory. It is much larger than the brick graph $B$, due to its large number of transitive edges. We implemented a dynamic programming algorithm that computes $A$ directly from $B$ without forming $\tilde{X}$.

This algorithm initializes $A:=B$, orders the nodes of $A$ so that it is lower-triangular, and iterates over its nodes from last to first; this way, when node $i$ is visited, its descendants have been visited already. When visiting node $i$, it identifies for every descendent $j$ of $i$ the parent nodes $k_{1},\ldots,k_{n}$ of $j$ that are also descendents of $i$:

$$i\to\ldots\to k_{1},\ldots,k_{n}\to j.$$

It iterates over these descendants in the forward direction – parents before children – and computes the total edge weight:

$$s_{ij}=A_{jk_{1}}+\cdots+A_{jk_{n}}.$$

If we can guarantee that $s_{ik_{1}}=s_{ik_{2}}=\cdots=s_{ik_{n}}=1$, then $s_{ij}$ is the path sum between $i$ and $j$ as well. We do so by adding an edge between $i$ and $j$ having weight $A_{ij}=1-s_{ij}$; if $A_{ij}\neq0$ already, then its weight is modified.

This algorithm guarantees that the resulting weighted DAG is one-summed because after visiting node $i$, the path sum between $i$ and any descendent $j$ is:

$$s_{ij}=A_{ij}+\sum_{\mathrm{parents}k \mathrm{of}j} s_{ik}A_{kj}$$

and recursively, $s_{ik}$ is 1 if $k$ is a descendent of $i$. Moreover, $s_{ij}$ is never modified after $i$ is visited, because all new edges involve non-descendants of $i$.

This algorithm could probably be made more efficient. Consider a simple path:

$$1\to2\to\cdots\to n$$

If the edge weights are all equal to one, then this graph is already one-summed, and this can be verified easily in time $O\left( n \right)$. However, the algorithm described here would run in $O\left( n^{2} \right)$ on this graph because it examines every ancestor-descendant pair. In practice, the brick graph is relatively shallow; most pairs of nodes are neither an ancestor nor a descendant of each other, and the runtime of this algorithm is smaller than that of the recombination step.

### 8. Genotype matrix operations on the ARG

We implemented memory efficient solvers for the ‘matmat’ operation

$$Y\leftarrow XB$$

as well as the ‘rmatmat’ operation

$$B\leftarrow X^{'}Y.$$

A simple approach to evaluate $XB=S\left( I-A \right)^{-1}MB$ involves the following three steps:

1. Compute $\tilde{B}=MB$, the matrix whose *i*th row is the sum of rows $B_{k}$ across variants $k$ that are assigned to node $i$. In practice, most rows of $MB$ are zero.
2. Solve ${\tilde{Y}=\left( I-A \right)}^{-1}\tilde{B}$. The *i*th row of $\tilde{Y}$ is the sum of rows $\tilde{B}_{j}$ among ancestors $j$ of $i$ in $A$.
3. The rows of $\tilde{Y}$ that correspond to sample nodes form the matrix $Y$.

In step (2), $A$ is a lower-triangular compressed sparse column (CSC) matrix, and $\tilde{Y}$ is computed via a ‘forward solve’ where columns $j$ are processed from first to last. At each iteration, row $j$ of $\tilde{B}$ is used to update its column neighbors’ rows: for all column neighbors $i$ of $j$,

$$\tilde{B}_{i}\leftarrow\tilde{B}_{i}+A_{ij}\tilde{B}_{j}.$$

This approach is fast but not memory efficient, as the result of every intermediate calculation is stored simultaneously in memory; most of these are discarded in step (3). An efficient approach leverages the approximately banded structure of the ARG, where ancestors of most nodes $i$ – in particular, ancestors of non-sample nodes – are likely to have indices close to $i$. The same is untrue of descendants because all nodes have samples as descendants. We assign to each node $i$ a *non-unique index*, $u\left( i \right)$, with the property that if $u\left( i \right)=u\left( j \right)$, $i<j$, then the row-neighbors (“parents”) of $j$ are greater than $i$:

$$\begin{aligned} i<k<j \#\left( * \right) \end{aligned}$$

for any $k$ such that $A_{kj}\neq0$.

To compute these indices, we initialize an array $d$ with the row-degree (number of parents) of each node. We initialize a stack of ‘available’ indices; when this stack is empty, it returns the next index that has not been used. We iterate over columns $j=n-1,\ldots,0$. We assign $u_{j}$ to be the first available index from the stack. For each column-neighbor (child) $i$ of the current node $j$, we decrement $d_{i}$. $i$ has already been visited, because the matrix is lower-triangular, so $u_{d_{i}}$ has already been assigned. If $d_{i}$ has reached 0, we push the index $u_{d_{i}}$ to the stack; some other node $j$ can be assigned $u_{d_{j}}=u_{d_{i}}$, as the condition (*) is satisfied.

We use this procedure to assign indices to non-sample and non-mutation nodes. Sample nodes and mutation nodes always get their own index, although two mutations in perfect LD may belong to the same mutation node (in which case they have the same index). In practice, the number of indices needed, $n'$, is much smaller than the number of nodes $n$, and not much different from the number of sample and mutation nodes.

We modify the simple forward solver to use the non-unique indices as follows. We initialize $\tilde{B}$ to be an all-zeros array of size $n^{'}\times k$ and add row $i$ of $B$ to row $u_{i}$ of $\tilde{B}$. We iterate over columns $j=0,\ldots,n-1$ of $A$. For each child $i$ of $j$, we update:

$$\tilde{B}_{u\left( i \right)}\leftarrow\tilde{B}_{u\left( i \right)}+A_{ij}\tilde{B}_{u\left( j \right)}.$$

After iterating over child nodes, $\tilde{B}_{u\left( j \right)}$ is reset to zero, unless $j$ has no children; this way, new values can be accumulated at the same index $u\left( j' \right)=u\left( j \right)$. Due to the choice of indices, we guarantee the row-neighbors (parents) $j^{'}>j$ are encountered after resetting $u\left( j \right)$. This procedure computes $\tilde{Y}$ in place, and because sample nodes do not have their values overwritten, we can extract the solution $Y$.

The backward solver uses parallel logic. To compute $B=X'Y$, we begin by assigning each row of $Y$ to the corresponding row of the $n^{'}\times k$ matrix $\tilde{Y}$. We iterate over columns $j=n-1,n-2,\ldots,0$. We first reset $\tilde{Y}_{u\left( j \right)}$ to zero, unless it has no children (i.e., it is a sample node). Then, for each child $i$ of $j$, we update:

$$\tilde{Y}_{u\left( j \right)} \leftarrow\tilde{Y}_{u\left( j \right)} +A_{ij}\tilde{Y}_{u\left( i \right)} .$$

This works for the same reason as the forward procedure. There are three differences between the forward and backward procedures: (1) the order in which columns are traversed; (2) which row is updated, the parent or the child; and (3) whether the reset-to-zero operation is performed before or after the row-update operation.

Our implementation uses one important optimization, which is the use of the ‘axpy’ BLAS subroutine to compute updates. This function increments an array $y$ by a scalar $\alpha$ (the edge weight) times an array $x$, and it runs significantly faster than a simple loop.

### 9. Computing the number of heterozygotes

The number of heterozygotes, $n_{het}$, is a useful quantity to compute as it allows the empirical variance to be computed without assuming HWE. We can write it as

$n_{het}=2n_{carriers}-n_{alleles}$,

where $n_{carriers}$ and $n_{alleles}$ are the number of carriers and alleles, respectively.

However, $n_{carriers}$ cannot be directly computed form the linear ARG. To address, this we can add individual nodes to the linear ARG such that each individual has incoming edges from its two haplotypes and then apply linearization. Since the resulting linear ARG is one-summed,

$n_{carriers}$is equivalent to the path sum from the all the individual nodes to the variant node. This can be computed with a leaf to root graph traversal with individual nodes inititalized to 1. Similarly, $n_{alleles}$ can be computed with another leaf to root graph traversal, but with all haplotype nodes initialized to 1. However, since the graph traversal procedure is additive, we can efficiently compute $n_{het}$ using a single graph traversal with individual nodes initialized to 2 and the haplotypes nodes to -1.

### Supplementary Tables and Figures

**Supplementary Table 1: Size in memory comparison.** Size in memory comparison of the linear ARG against CSC matrix and GRG on the UK Biobank 200k across all variants on chr 1, 11, 21.

| **Data structure** | **Size in memory (GB)** |
| --- | --- |
| Sparse matrix (CSC) | 447.5 |
| GRG | 20.6 |
| Linear ARG | 8.2 |

**Supplementary Table 2: UK Biobank 200k chr 1, 11, 21 disk size comparison.** Disk size comparison of the linear ARG against four other data representations across all variants on chr 1, 11, 21.

| **Data structure** | **Disk size (GB)** |
| --- | --- |
| vcf.gz | 333.7 |
| .pgen | 55.8 |
| XSI | 11.1 |
| GRG | 9.3 |
| Linear ARG | 3.8 |

**Supplementary Table 3: Linear ARG inference costs.** Cost to infer linear ARGs on UKB-RAP. UKB 200k chr1-22 was run on low priority while the other two datasets were run on high priority. All of Us does not provide costs for each job, so only total cost is provided.

|  | UKB 200k chr1-22 (£) | UKB 200k MAF>0.01 chr1-22 (£) | UKB 500k chr21 (£) | All of Us chr21 ($) |
| --- | --- | --- | --- | --- |
| step1 | 212.81 | 592.45 | 36.88 | NA |
| step2 | 671.86 | 892.58 | 49.17 | NA |
| step3 | 287.29 | 165.93 | 19.18 | NA |
| step4 | 493.61 | 218.85 | NA | NA |
| total | 1,665.57 | 1,869.81 | 105.23 | 141.88 |

**Supplementary Table 4: Phenotypes and covariates.** Identifiers and names of the 89 phenotypes and 42 covariates used in the GWAS benchmark.

**Supplementary Table 5: UKB 200k linear ARG sizes by chromosome.** Disk size, size in memory, and number of nonzeros of the UKB 200k linear ARGs.

**Supplementary Table 6: UKB 200k zipped VCF and .pgen files sizes by chromosome.** Disk size of vcf.gz, .pgen, .psam, and .pvar for the UKB 200k data without variant filtering and MAF threshold of 0.01.

**Supplementary Table 7: Size benchmark of other data representations on UKB 200k chr 1, 11, 21.** Disk size of GRG and XSI and size in memory of GRG and Scipy CSC matrix by chromosome.

**Supplementary Table 8: Linear ARG compression on msprime-simulated data.** Mean and standard deviation of the number of nonzeros in the genotype matrix, inferred linear ARG, and data-generating ARG across 100 simulations.

**Supplementary Table 9: Robustness of the linear ARG compression ratio in simulation.** Mean and standard deviation of brick graph and linear ARG nonzero ratio after simulating back mutations, recurrent mutations, genotyping errors, and out-of-order errors across 100 simulations.

**Supplementary Table 10: GWAS runtime and peak memory usage benchmark.** Runtime and peak memory usage of PLINK 2.0 and kodama for performing a GWAS on the UKB 200k dataset across 89 traits with 42 covariates.

**Supplementary Table 11: PLINK 2.0 and kodama GWAS concordance.** Correlation of beta, standard error, and z-score between PLINK 2.0 and kodama across variants and stratified by trait.

**Supplementary Table 12: Matrix multiplication benchmark.** Load time, matrix multiplication time, and peak memory usage for the linear ARG, GRG, and XSI data structures.

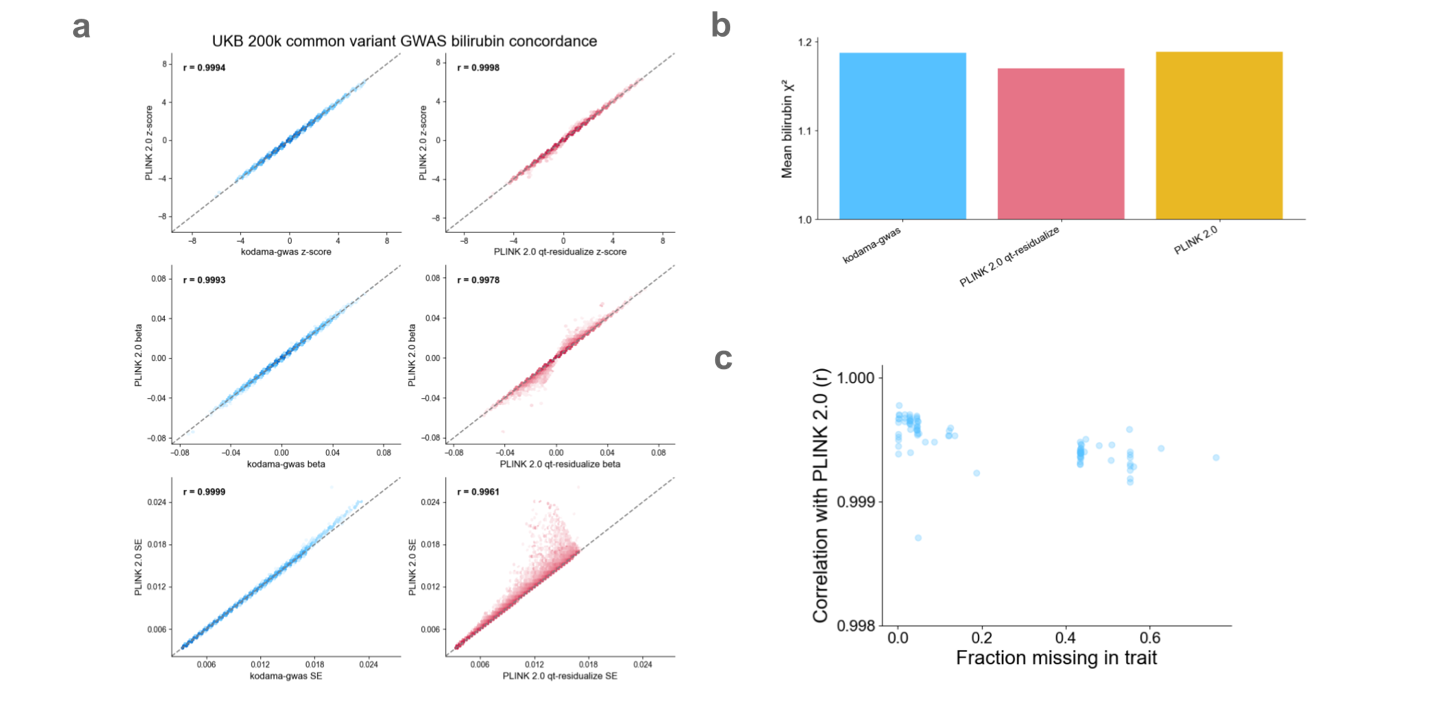

**Supplementary Figure 1: UKB 200k Common variant GWAS benchmark. a,** Concordance of z-scores, betas, and standard errors compared to PLINK 2.0 for bilirubin (0.18 missingness) on chromosome 11. **b,** Mean chi-squared value for bilirubin on chromosome 11 across methods. **c,** Correlation between PLINK 2.0 and linear ARG z-scores for each trait across chromosomes 1-22.

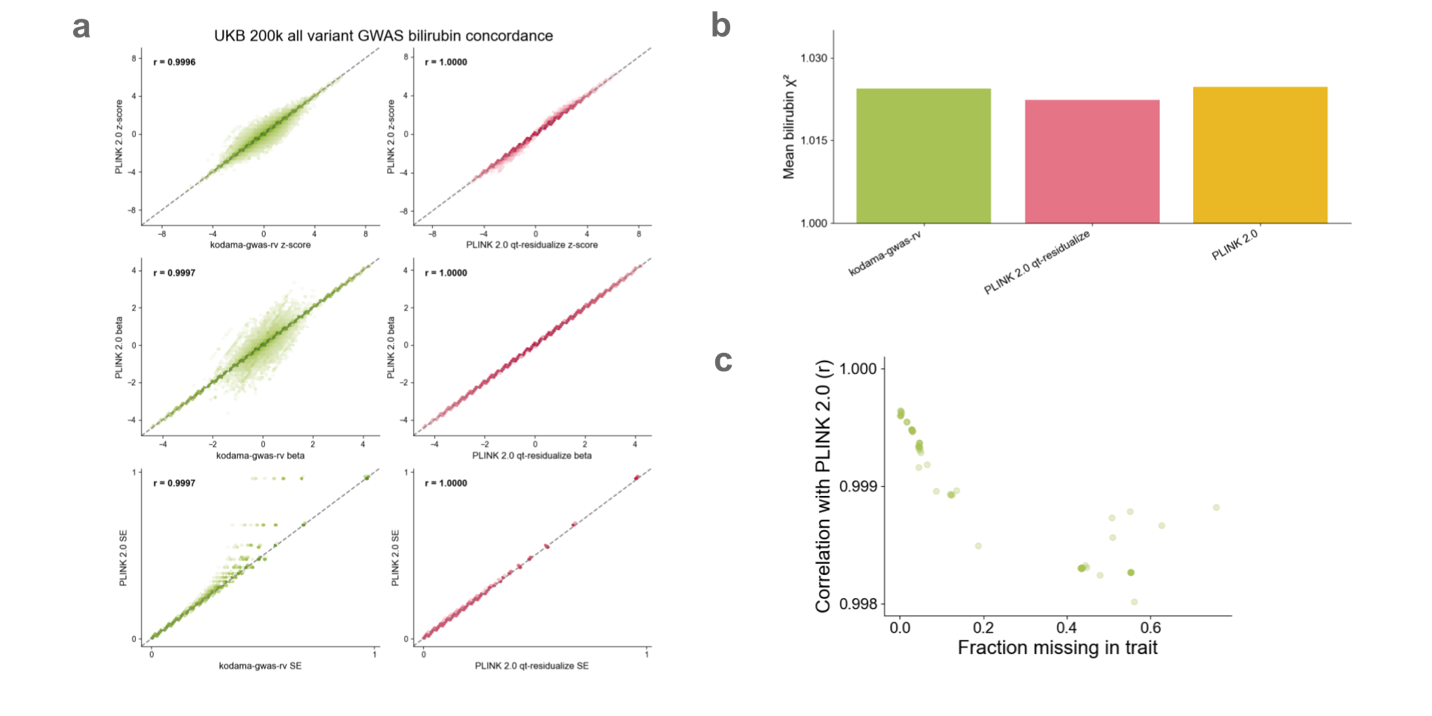

**Supplementary Figure 2: UKB 200k all variant chr 1, 11, 21 GWAS benchmark. a,** Concordance of z-scores, betas, and standard errors compared to PLINK 2.0 for bilirubin (0.18 missingness) on chromosome 11. **b,** Mean chi-squared value for bilirubin on chromosome 11 across methods. **c,** Correlation between PLINK 2.0 and linear ARG z-scores for each trait on chromosome 11.

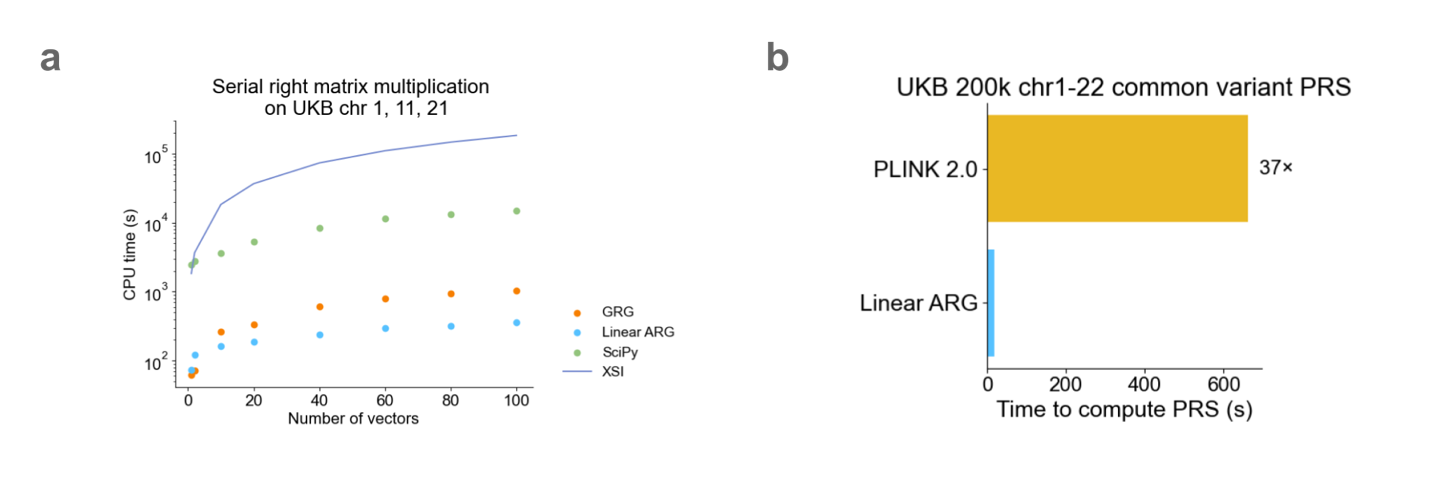

**Supplementary Figure 3: Matrix multiplication runtime benchmark. a,** Composite time to perform matrix multiplication (including load time, if applicable) on matrices of sizes 1 to 100. Since XSI has only implemented matrix-vector multiplication, the purple line represents the time it takes to perform this matrix-vector multipication multiplied by the size of the matrix. **b,** PRS runtime comparison between the linear ARG and PLINK 2.0 on the UKB 200k common variant dataset on the for a single trait.
